# *In vivo* neural activity of electrosensory pyramidal cells: Biophysical characterization and phenomenological modeling

**DOI:** 10.1101/2025.05.30.656684

**Authors:** Amin Akhshi, Michael G. Metzen, Maurice J. Chacron, Anmar Khadra

**Affiliations:** Department of Physiology, McGill University, Montreal, QC, H3G 1Y6, Canada

**Keywords:** Pyramidal cells, burst firing, electrosensory system, conductance-based biophysical model, modified Hindmarsh-Rose model

## Abstract

Burst firing is an important property of neuronal activity, thought to enhance sensory encoding. While previous studies show significant differences in burst firing between *in vivo* and *in vitro* conditions, how burst firing contributes to neural coding *in vivo* and how it is modulated by underlying biophysical mechanisms when neurons are under active synaptic bombardments remains poorly understood. Here, we combined intracellular recordings and computational modeling to investigate how cellular and synaptic mechanisms can explain the *in vivo* firing activity of electrosensory lateral line lobe (ELL) pyramidal cells in *Apteronotus leptorhynchus*. We developed a biophysically detailed compartmental model incorporating voltage-gated currents, NMDA receptor-mediated Ca^2+^ influx, Ca^2+^-activated SK channels, Ca^2+^ handling, and stochastic synaptic inputs to reproduce *in vivo* firing activities of ELL pyramidal cells. Specifically, using bifurcation analysis, we identified dynamical transitions between quiescent, tonic, and bursting regimes, governed by interactions among SK conductance, NMDA receptor activation, and applied current. Model parameters were optimized against *in vivo* data, accurately reproducing action potential waveforms and temporal dynamics, including characteristic bimodal interspike interval distributions reflecting intra- and inter-burst intervals. We further developed a modified Hindmarsh-Rose model incorporating dual adaptation variables and stochastic noise. This simplified phenomenological model successfully captured burst firings comparable to those observed in the biophysical model and recorded data, while replicating diverse firing patterns observed across the population. Finally, parameter sensitivity analysis revealed slow adaptation dynamics and noise intensity as key determinants of spiking variability within cells. Overall, our modeling results demonstrate that *in vivo* bursting arises from synergistic interactions between intrinsic conductances (e.g., NMDA-SK coupling), Ca^2+^ mobilization, and synaptic stochasticity, offering a potential reconciliation for discrepancies with *in vitro* firing activity. The models provide mechanistic insights into how background synaptic activity modulates burst firing and validate simplified frameworks for studying population-level dynamics.

## Introduction

Understanding how neurons in sensory systems process incoming information to drive perception and behavior remains a fundamental challenge in neuroscience. In particular, this understanding is complicated by the fact that neural populations display complex spiking activity patterns, even within a given neural class (Bannister and Larkman, 1995a, b; Gjorgjieva et al., 2016), which are thought to enhance the encoding of stimulus-relevant information (Krahe and Gabbiani, 2004; Tan et al., 2014; Horrocks et al., 2024). Growing evidence, however, suggests that such firing patterns can differ significantly between *in vivo* and *in vitro* conditions (Bernander et al., 1991; Destexhe and Paré, 1999; Destexhe et al., 2003; Prescott et al., 2008; Akerberg and Chacron, 2011; Metzen et al., 2016). Indeed, in the intact behaving brain, neurons collectively receive high levels of background synaptic activity that substantially alter their membrane properties and modulate their integrative function (i.e., firing activity patterns) (Destexhe et al., 2001; Destexhe et al., 2003; Haider and McCormick, 2009; Reinhold et al., 2015), in contrast to *in vitro* conditions where circuit effects are largely absent (Destexhe et al., 2003; Prescott et al., 2008; Wei et al., 2023). Specifically, such synaptic bombardment *in vivo* introduces membrane potential fluctuations that bring neurons closer to the firing threshold, thereby enhancing responsiveness and improving temporal precision (Fellous et al., 2003; Tan et al., 2014; Pala and Petersen, 2015; Pala and Petersen, 2018; Akhshi et al., 2023). These changes are potentially functionally relevant as they are thought to improve information coding efficiency due to changes in the timescale of neural responses and synchronization characteristics (Destexhe et al., 2003; Mejias and Longtin, 2012; Metzen and Chacron, 2023; Koch et al., 2025). Together, these effects underscore the critical importance of understanding how *in vivo* conditions influence neuronal dynamics, a topic that remains to be systematically explored.

One of the most prominent examples of a neural system exhibiting *in vivo* vs. *in vitro* discrepancies is found in pyramidal cells within the electrosensory lateral line lobe (ELL) of weakly electric fish (Toporikova and Chacron, 2009; Avila-Akerberg and Chacron, 2011; Deemyad et al., 2011; Metzen et al., 2016). The electrosensory system of these fish offers a well-characterized neural circuit, making it advantageous for studying how *in vivo* conditions affect neural activity patterns (Bell and Maler, 2005; Chacron et al., 2011; Krahe and Maler, 2014; Metzen and Chacron, 2019). These animals generate a continuous electric organ discharge (EOD) used for navigation and communication, and detect perturbations of this signal via electroreceptor afferents (EAs) distributed across the skin, which relay sensory input to pyramidal cells within the ELL (Bastian, 1986; Turner et al., 1999) that project to the midbrain (Sas and Maler, 1983, 1987; Huang et al., 2019; Metzen and Chacron, 2019, 2021, 2023). A defining feature of ELL pyramidal cells is their ability to fire action potentials in the form of bursts (i.e., clusters of spikes followed by quiescence). *In vitro* studies of ELL pyramidal cells have revealed that burst firing relies on a somato-dendritic interplay involving the backpropagation of action potentials from the soma to the apical dendrites (Lemon and Turner, 2000; Doiron et al., 2002). This process is associated with a progressive increase in the amplitude of depolarization afterpotentials (DAPs) at the soma and shortening interspike intervals (ISIs) until the burst terminates with dendritic failure and a subsequent large burst afterhyperpolarization (AHP) (Turner et al., 1994; Doiron et al., 2001; Fernandez et al., 2005). The *in vivo* recordings, on the other hand, show markedly different firing patterns, typically consisting of only 2–3 spikes per burst on average, with no evidence of a somatic DAP or burst AHP (Bastian and Nguyenkim, 2001; Toporikova and Chacron, 2009; Deemyad et al., 2011; Deemyad et al., 2013). Notably, pharmacological blockade of intracellular Ca^2+^ using BAPTA restores *in vitro*-like burst firing patterns *in vivo* (Ellis et al., 2007b; Krahe et al., 2008; Mehaffey et al., 2008b; Toporikova and Chacron, 2009), implicating the important role of Ca^2+^ dynamics and Ca^2+^-activated SK channels as critical modulators of burst dynamics under *in vivo* condition. The disparity between the *in vivo* and *in vitro* recordings suggests that detailed computational models are required to account not only for additional intrinsic ionic conductances expressed *in vivo* but also for extrinsic factors that influence spiking activity.

In this study, we combined intracellular *in vivo* recordings of ELL pyramidal cells in *Apteronotus leptorhynchus* with computational modeling to investigate how biophysical mechanisms such as Ca^2+^ dynamics, SK and NMDA receptor currents, as well as stochastic synaptic inputs, can explain discrepancies between neuronal firing activities seen *in vitro* and *in vivo*. First, we built a detailed biophysical model and constrained its parameter regimes to accurately reproduce both the waveform and the temporal dynamics of action potentials.

We subsequently employed bifurcation analysis to systematically examine the influence of key parameters on the critical transitions between quiescent, tonic, and burst firing regimes. Next, we developed a phenomenological model based on the Hindmarsh-Rose (HR) formalism and demonstrate that the modified HR model accurately reproduces the spiking activity of ELL pyramidal cells, providing a computationally efficient framework with significant advantages for modeling population coding in neural circuits.

## Results

The goal of this study was to use computational modeling to investigate how cellular and synaptic mechanisms can explain experimentally observed differences in firing activity of ELL pyramidal cells *in vitro* and *in vivo*. ELL pyramidal cells receive direct ascending inputs from electroreceptor afferents (EAs; Fig. 1A, black arrows) as well as substantial descending feedback (Fig. 1A, orange arrows) (Sas and Maler, 1983, 1987). These inputs regulate firing activity through a combination of synaptic mechanisms involving both glutamatergic excitation (e.g., mediated in part by NMDA receptors) (Berman et al., 1997; Huang et al., 2019; Metzen and Chacron, 2023) and feedback-activated inhibition (Berman and Maler, 1998; Lewis and Maler, 2002). We performed single-unit intracellular recordings *in vivo* from *n* = 20 pyramidal cells in the ELL across *N* = 3 fish in the absence of stimulation (Fig. 1A, left; see Methods for details). Figure 1B shows the membrane potential trace for a representative cell (Fig. 1B, left; 5 s duration) displaying spiking activity characterized by periods of rapid, clustered spiking, consistent with burst firing. Notably, and in contrast to firing patterns observed *in vitro* (Lemon and Turner, 2000; Doiron et al., 2001; Rashid et al., 2001; Doiron et al., 2002), these *in vivo* bursts consisted of fewer spikes and lacked prominent DAPs and distinct burst-specific AHPs. Consistent with the clustered spiking observed in the voltage trace (Fig. 1B left), the interspike interval (ISI) distribution for this neuron was bimodal (Hartigan’s dip test, p-value = 0.00402) (Fig. 1B, right), with a prominent peak at short ISIs (< ∼12 ms) corresponding to the intra-burst interval and a peak at larger values (∼100 ms) corresponding to the inter-burst interval. To quantify burst firing across the population, we defined bursts based on a threshold set at the local minimum between these two ISI modes to separate spike trains into burst and isolated spikes for each cell (Oswald et al., 2004); a burst was identified as any sequence of two or more spikes in which all consecutive ISIs fell below this threshold (Supplementary Fig. 1A; see Methods). Unlike approaches that treat bursts as unitary events, we explicitly accounted for burst structure, including the number of spikes within each burst, and computed the fraction of spikes identified as belonging to bursts.

**Figure 1.**
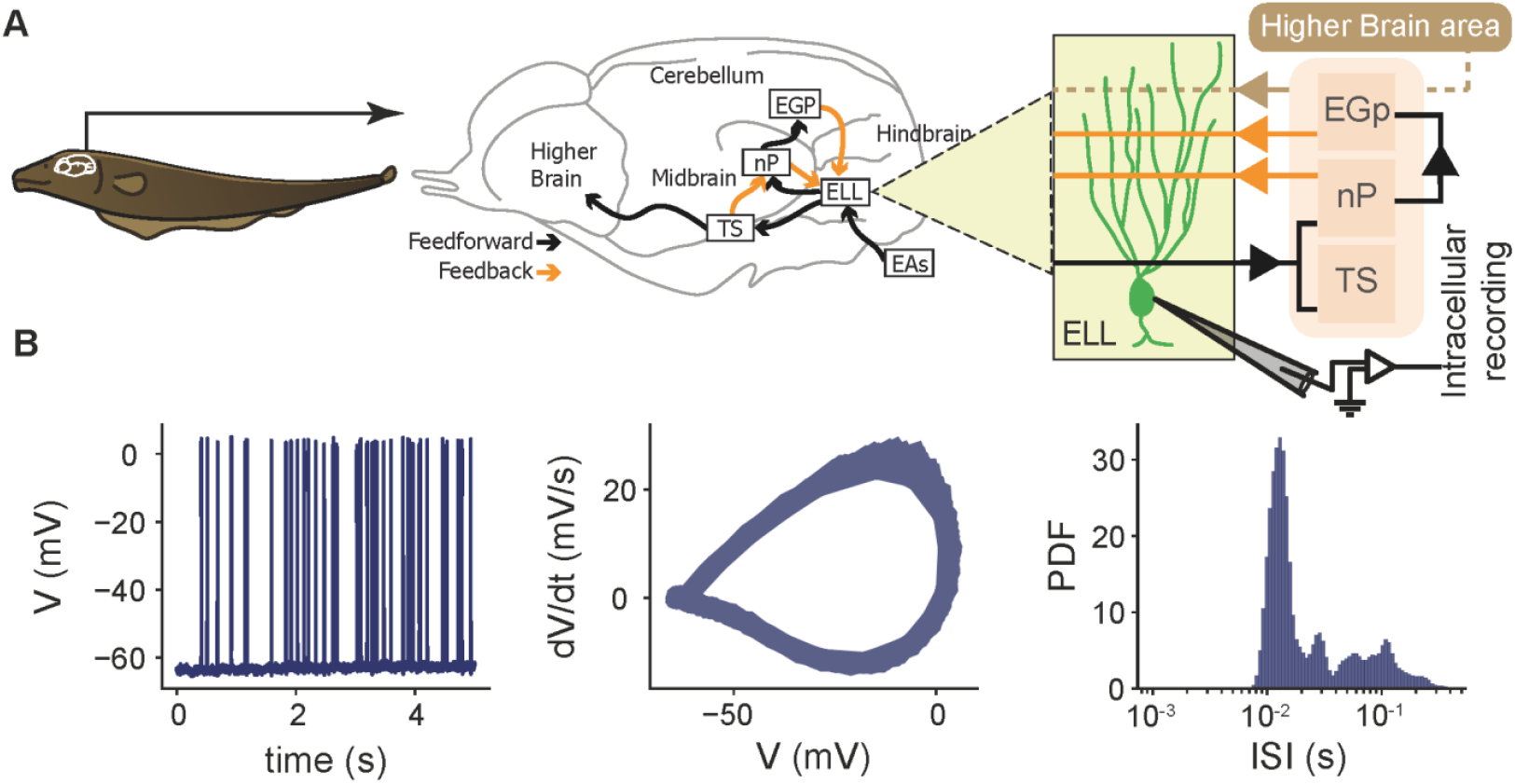
Circuit and single cell electrical properties of electrosensory lateral line lobe (ELL) pyramidal cells. **(A)** Schematic of the neural circuitry underlying the electrosensory processing in the weakly electric fish (left). Brain circuitry (middle) showing how the ELL receives feedforward input from electrosensory afferents (EAs) and feedback signals from higher brain areas, including the eminentia granularis posterior (EGp), nucleus praeminentialis (nP), and torus semicircularis (TS). Intracellular recordings were obtained from pyramidal neurons within the ELL to investigate their electrical activity (right). **(B)** Intracellular membrane potential trace recording for 5s (left), phase plane plot of action potential cycles in the voltage trace (middle), and interspike interval (ISI) distribution (right) for a representative electrophysiological data from an ELL pyramidal cell.

Consistent with previous studies (Bastian and Nguyenkim, 2001; Bastian et al., 2002; Krahe et al., 2008; Akhshi et al., 2023), we observed that the mean firing rate across the recorded population ranged between 5-40 Hz in the absence of stimulation (Supplementary Fig. 1B, left), and that spike train statistics including the tendency for burst firing varied considerably between cells (Supplementary Fig. 1B, right).

### A biophysical model of ELL pyramidal cells accurately reproduces the in vivo action potential characteristics

To gain understanding as to how different mechanisms can account for differences between *in vivo* vs *in vitro* bursting dynamics in ELL pyramidal cells, we developed a new conductance-based biophysical model, partially inspired by previous studies (Doiron et al., 2002; Toporikova and Chacron, 2009), that incorporates the flux balance model of Ca^2+^ mobilization (Fig. 2A; see Methods). As done previously (Doiron et al., 2002), the intrinsic bursting patterns of ELL pyramidal cells were captured by representing the soma and dendrites as two coupled isopotential compartments linked by a resistive current scaled by the soma-to-dendrite area ratio (*κ*), with each compartment containing distinct ionic conductances (Fig. 2A; see Methods). The two compartments included fast Na^+^ (*I*_*Na,S*_, *I*_*Na,D*_, respectively) and delayed rectifier K^+^ (*I*_*Dr,S*_, *I*_*Dr,D*_, respectively) currents, both responsible for generating action potentials. The dendritic compartment, however, included additionally Ca^2+^-activated small conductance K^+^ current (*I*_*SK*_) that hyperpolarizes the membrane voltage in response to intracellular Ca^2+^ increases and helps modulate burst termination, as well as a Ca^2+^ current due to NMDA receptors (*I*_*NMDA*_), the primary source of Ca^2+^ influx mediating dendritic depolarization (Fig. 2A; dendrite). The kinetics of the NMDA receptors was based on a Markov chain model (Destexhe et al., 1994) consisting of three closed states (*C*_0_,*C*_1_,*C*_2_), one conducting open state (*O*) and one desensitized state (*D*) (see Methods), allowing the receptors to generate currents that are voltage-dependent due to Mg^2+^ block ([*Mg*^2+^]_*o*_). During depolarization, the magnesium block is relieved, allowing Ca^2+^ influx into the cell. This creates a regenerative depolarizing current that sustains dendritic spikes and amplifies burst generation. The flux-balance model for Ca^2+^ mobilization in the dendritic compartment was based on the Li-Rinzel formalism, accounting for Ca^2+^ entry through NMDA receptors, Ca^2+^ buffering in the cytosol to keep it at low concentration, IP3 receptor flux via Ca^2+^-induced Ca^2+^-release mechanism from the endoplasmic reticulum (ER), SERCA flux that pumps Ca^2+^ back into the ER for sequestration, PMCA flux that pumps Ca^2+^ out of the cell and leak fluxes through plasma and ER membranes (Li and Rinzel, 1994; Oprea et al., 2022). The resulting model generates cytosolic Ca^2+^ transients that activate *I*_*SK*_, contributing to the termination of bursts by inducing afterhyperpolarization (Fig. 2A; right). The ER Ca^2+^ stores, on the other hand, can deplete and recover, introducing temporal variability in burst initiation and duration. To account for the contribution of synaptic inputs to burst generation, we included a synaptic input current (*I*_*syn*_) in the dendritic compartment to represent the stochastic background excitatory and inhibitory pre-synaptic activity (Manwani and Koch, 1999a; Brake et al., 2024; Koch et al., 2025). Under stochastic synaptic input, the model generated voltage dynamics and Ca^2+^ transients that are very similar to those observed *in vivo* (Supplementary. Fig. 2).

**Figure 2.**
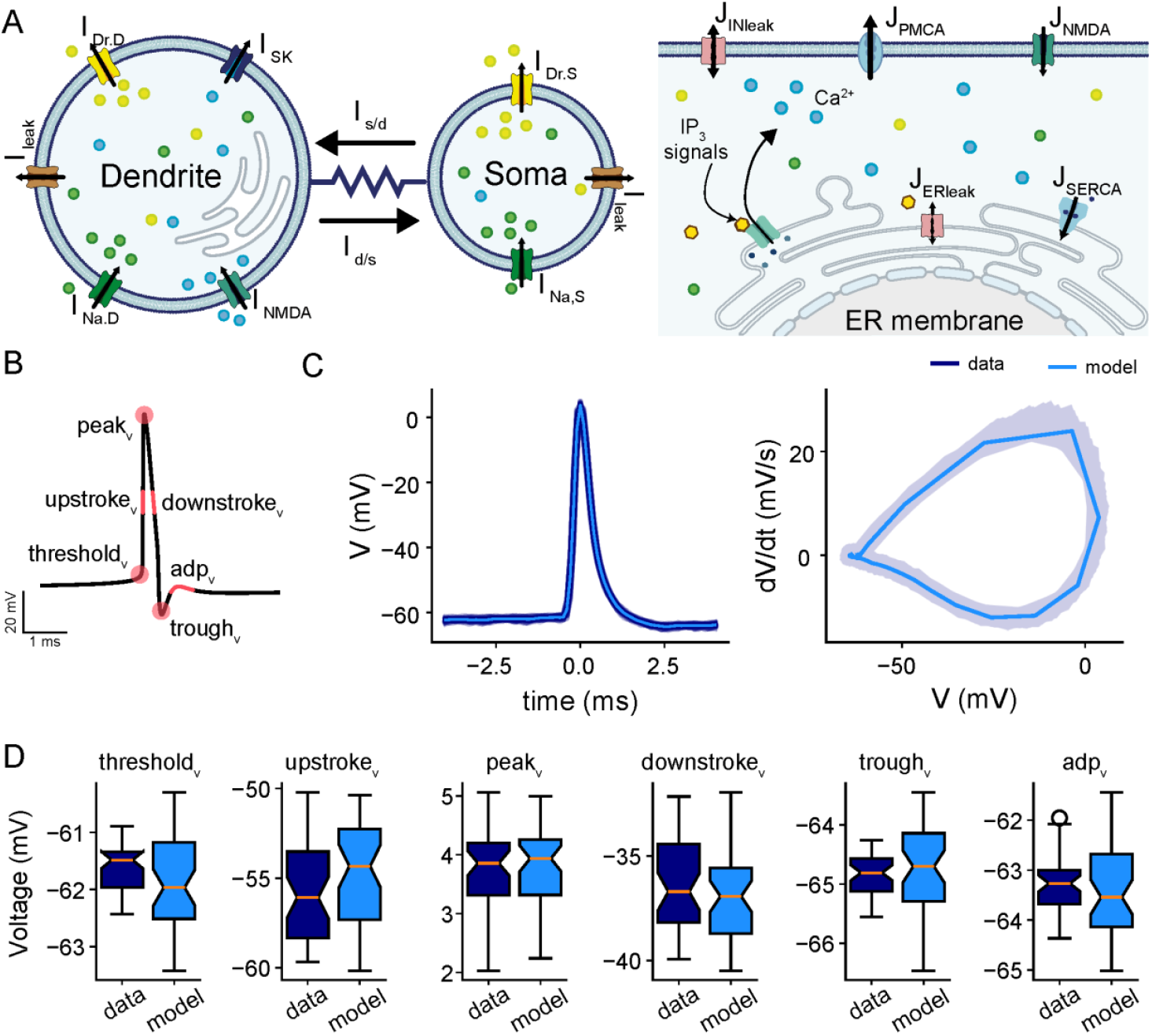
Comparison of *electrophysiological features between intracellular recording of a given ELL pyramidal cell and a model simulation*. **(A)** Schematic of the model showcasing the two-compartments of the cell: soma and dendrite (left), and highlighting the various ionic currents expressed in each compartment, including fast Na^+^ (*I*_*Na,i*_), delayed rectifier K^+^ (*I*_*K,i*_), small-conductance K^+^ (*I*_*SK*_), NMDA (*I*_*NMDA*_) and leak (*I*_*leak*_) currents in the soma (*i* = *S*) and dendrite (*i* = *D*). The kinetics of NMDA receptors are governed by a Markov model (gray box) consisting of three closed states (*C*_0_,*C*_1_,*C*_2_), one conducting open state (*O*) and one desensitized state (*D*). The schematic of Ca^2+^ dynamics within the dendritic compartment following the flux-balance model is also shown (right). It displays all fluxes involved in regulating Ca^2+^ mobilization across plasma and ER membranes in the dendritic compartment. These fluxes include Ca^2+^ entry through NMDA receptors (*J*_*NMDAR*_) and IP3 receptors (*J*_*IP*3*R*_), Ca^2^+ efflux through SERCA (*J*_*SERCA*_) and PMCA (*J*_*PMCA*_) pumps, and leak across both membranes. **(B)** Schematic of an action potential with the extracted electrophysiological features (highlighted in pink) for model fitting; that includes from left to right: spike threshold (threshold_v_), midpoint of the upstroke phase (upstroke_v_), peak amplitudes (peak_v_), midpoint of the downstroke phase (downstroke_v_), trough amplitudes (rough_v_) and amplitude of afterdepolarization potential (adp_v_). **(C)** Extracted action potentials (left) and action potential cycles (right) obtained from one recorded trace (5 sec; shaded dark blue) and the average action potential and average action potential cycle of all spikes obtained from a single model simulation overlaid on top (5 sec; light blue). **(D)** Box plots of electrophysiological features obtained from all action potentials extracted from the experimental recording and modeling simulation in C. Panels from left to right: threshold_v (_, upstroke_v_, peak_v_, downstroke_v_, rough_v_, adp_v_. Two-sample t-tests were performed for each feature, with the following p-values: p_threshold_= 0.054, p_upstroke_ = 0.152, p_peak_ = 0.694, p_downstroke_ = 0.395, p_trough_ = 0.502, p_adp_ = 0.283.

To assess the performance of our model and identify the physiologically relevant parameter ranges *in vivo*, we fitted the model to physiological features of action potentials extracted from intracellular recordings (see Methods). Feature extraction followed the Allen Institute’s protocols for electrophysiological data processing, which include features such as voltage threshold for spiking, peak amplitudes, trough amplitudes, voltage at midpoint of the upstroke phase, voltage at midpoint of the downstroke phase, and amplitude of afterdepolarization potential (highlighted in pink; Fig. 2B). To isolate individual action potentials, we extracted segments of the voltage time series centered around the peak amplitudes within an 8 ms window (i.e., 4 ms before and after the peak; see Methods). A comparison of extracted features from experimental recordings (dark blue) and model simulations (light blue) revealed a close match, as evidenced by the overlaid simulated spikes on recorded spikes for a representative ELL pyramidal cell (Fig. 2C); the model accurately reproduces the shape and defining characteristics of action potentials. This agreement was further quantified by statistical comparisons of each feature between the experimental data and model simulations using two-sample t-tests, confirming no significant differences for any of the examined features (Fig. 2D; p_threshold_= 0.054, p_upstroke_ = 0.152, p_peak_ = 0.694, p_downstroke_ = 0.395, p_trough_ = 0.502, p_adp_ = 0.283). Furthermore, to assess the model’s performance across the population, we compared the mean value of each feature between the model and experimental data across all recorded cells (Supplementary Fig. 3; linear fit: r_threshold_ = 0.96, p_threshold_ = 2.5 × 10^−12^; r_upstroke_ = 0.98, p_upstroke_ = 1.1 × 10^−14^; r_peak_ = 0.99, p_peak_ = 4.9 × 10^−17^; r_downstroke_ = 0.99, p_downstroke_ = 4.7 × 10^−17^; r_trough_ = 0.98, p_trough_ = 9.7 × 10^−16^; r_adp_ = 0.98, p_adp_ = 2.1 × 10^−14^). The results showed points clustering tightly around the identity line, indicating strong agreement and demonstrating that the model accurately captures the average feature characteristics across the cell population. Overall, these results show that the biophysical two-compartment model effectively captures the physiological characteristics of *in vivo* action potentials in ELL pyramidal cells.

### The model captures burst dynamics and spiking variabilities observed across the ELL pyramidal cell population in vivo

Next, we assessed the ability of the model to reproduce the temporal firing dynamics of ELL pyramidal cells *in vivo* by comparing key spiking activity features between the data and model simulations obtained after parameter optimization (see Methods). Figure 3A compares representative voltage traces (top panels) and corresponding ISI distributions (bottom panels) between the example recording data (left) and the model simulation fitted to that data (right). Both traces exhibit qualitatively similar spiking patterns, including sequences of bursts interspersed with isolated spiking events. The corresponding ISI distributions show a close alignment (Kolmogorov-Smirnov test: D = 0.335, p=0.635), accurately preserving the characteristic bimodality observed in the data. In both distributions, the left mode primarily corresponds to fast spiking events within bursts (intraburst spikes), while the second mode at longer intervals reflects the timing between bursts or isolated spikes. Furthermore, raster plots comparing data and model outputs demonstrate that the model accurately reproduces the qualitative structure of these bursts and isolated spikes (Fig. 3B). Interestingly, the most commonly observed burst event patterns in both the data and the model included a few spikes within each burst, consistent with previous studies (Bastian and Nguyenkim, 2001; Deemyad et al., 2013). The close match in both ISI distributions and spike timing structure highlights the model’s ability to capture key temporal dynamics of both burst firing and regular spiking activity *in vivo*.

**Figure 4.**
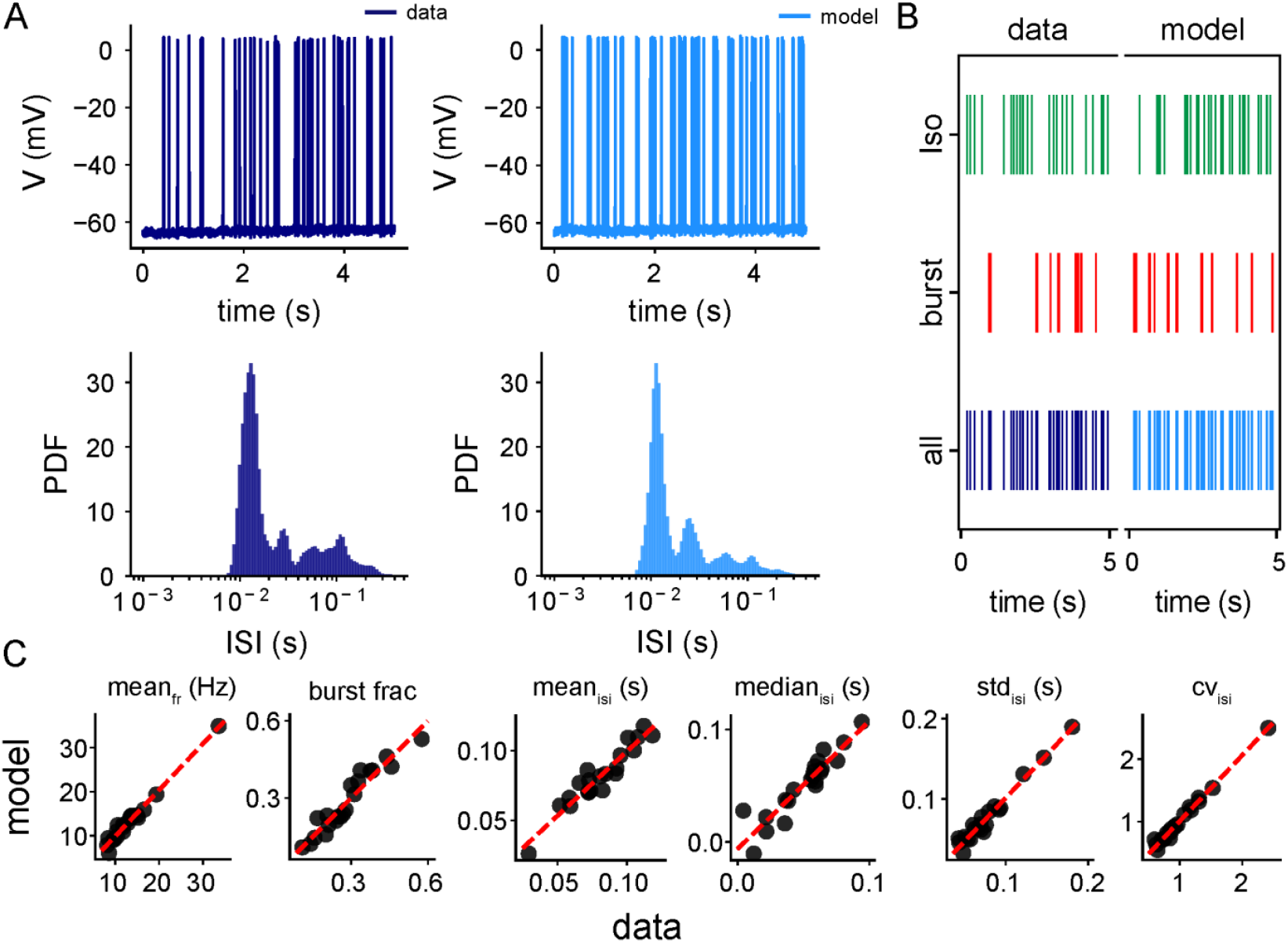
Comparison of spike train features between intracellular recordings of a population of ELL pyramidal cells and their fitted model simulations. **(A)** Membrane potential traces (top) and interspike interval (ISI) distribution (bottom) of a representative ELL pyramidal cell (dark blue, left) and its fitted model simulations (light blue, right). The recorded and simulated traces are 5 seconds long, showing sequences of bursts interspersed with isolated spikes and quiescent periods. **(B)** Raster spike train plots from same recording in A (left) and their corresponding fitted simulations (right), separated into isolated spikes (green), burst spikes (red), and all spikes combined (blue). The model replicates the proportions and temporal organization of burst and isolated spikes observed in the recordings. **(C)** Scatter plots comparing key spike train features between recordings and simulations for all ELL pyramidal cells (*n* = 20). Panels from left to right: mean firing rate (mean_fr_; r = 0.99, p=2.4 × 10^−16^), burst fraction (burst frac; r = 0.96, p=1.1 × 10^−11^), mean (mean_isi_; r = 0.95, p=3.5 × 10^−11^), median (median_isi_; r = 0.93, p=1.7 × 10^−9^), standard deviation (std_isi_; r = 0.98, p=1.7 × 10^−15^) and the coefficient of variation (CV_isi_; r = 0.99, p=1.4 × 10^−19^) of ISIs, respectively.

To quantify the overall performance of the model in reproducing the temporal firing activities of all recorded ELL pyramidal cells (*n* = 20), we systematically compared key spike train features generated by these cells to those generated by model simulations; that included: the mean firing rate, burst fraction, mean and median of ISI, the standard deviation of ISI, and the coefficient of variation for each recorded ELL pyramidal cell compared with the representative model neuron obtained by the best fit for that cell (Fig. 3C). Burst fractions for model simulations were obtained using the same method as for the data (see Methods). The pairwise comparison of all features showed a strong linear correspondence between experimental and simulated values across the population. All points clustered closely around the identity line, indicating that the model accurately reproduced cell-specific variability in firing activity (Fig. 3C; linear fit: mean firing rate (r = 0.99, p=2.4 × 10^−16^), burst fraction (r = 0.96, p=1.1 × 10^−11^), mean (r = 0.95, p=3.5 × 10^−11^), median (r = 0.93, p=1.7 × 10^−9^), standard deviation (r = 0.98, p=1.7 × 10^−15^), and coefficient of variation (r = 0.99, p=1.4 × 10^−19^) of ISI distributions, respectively). This further indicates that the temporal structure of spiking activities *in vivo* across the recorded population is well-preserved by the model.

Taken together, these results highlight that the biophysical model accurately reproduces variability in the burst firing dynamics of ELL pyramidal cells *in vivo*.

### Bifurcation analysis of biophysical model elucidate how intrinsic mechanisms affect spiking activity

We next investigated the deterministic dynamics of the model by exploring the effects of varying key model parameters including the applied current (*I*_*app*_) and the maximal conductances of *I*_*SK*_ and *I*_*NMDA*_ (*g*_*SK*_ and *g*_*NMDA*_, respectively) to understand how their effect on spiking activity. Our one-parameter bifurcation analysis revealed that varying *I*_*app*_ produces four distinct firing regimes: two quiescent (i.e., no action potential firing), one tonic firing (i.e., periodic firing of action potentials) and one ghostbursting regime; these regimes are separated by different types of bifurcation points (Fig. 4A). For low values of *I*_*app*_, the model exhibits three branches of equilibria: a stable one (bottom branch; solid orange line) corresponding to the quiescent state of the cell, and two unstable ones (middle and top branches; dashed black lines). Together, the three branches form an s-shaped manifold with two saddle nodes: one to the right (labeled SN1) where the bottom two branches merge, forming the right boundary of the left quiescent regime, and one to the left (labeled SN2) where the top two branches merge. The presence of three coexisting steady states to the left of SN1 enables the model to generate (among other things; see below) single spikes in response to prolonged suprathreshold constant current injections (Supplementary Fig. 4A). These spikes arise from type IV excitability (Mitry et al., 2020; MacKay et al., 2021), triggered when trajectories cross the threshold defined by the stable manifold of the saddle fixed points (the middle equilibria).

**Figure 4.**
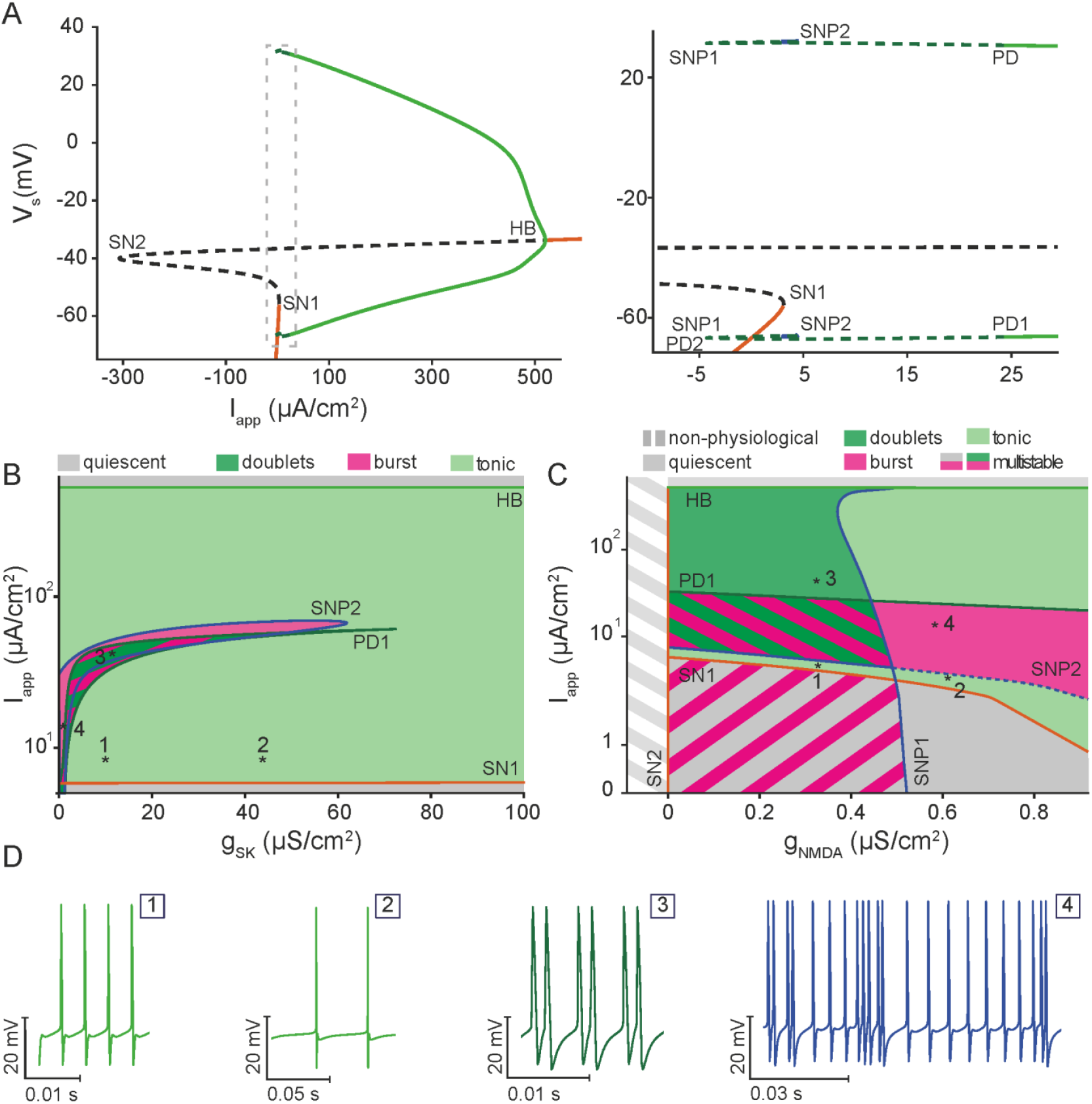
Bifurcation analysis of the deterministic biophysical model reveals the full dynamical behavior of ELL pyramidal cells. **(A)** One-parameter bifurcation diagram of the voltage variable (*V*) of the biophysical model with respect to the applied current (*I*_*app*_), illustrating the various regimes of behavior corresponding to branches of stable equilibria (orange line), branches of unstable equilibria (dashed black line), envelopes of periodic orbits (green lines) and envelopes of bursting orbits (dashed green line). Within the range of *I*_*app*_ considered, the model undergoes several types of bifurcation points, including 2 saddle-node bifurcations (SN1, SN2), 1 Hopf bifurcations (HB), 2 saddle-node of periodics bifurcations (SNP1, SNP2) and 2 period-doubling bifurcations (PD1, PD2), some of which define the boundaries of the various regimes of behavior identified. The inset provides magnified views of the area enclosed by the bounding boxes in the main figure. **(B)** The two-parameter bifurcation of the biophysical model with respect to *I*_*app*_ and the maximum conductance of SK channels (*g*_*sk*_), highlighting the various regions of behavior that the model possesses, including quiescence, tonic firing and doublet/chaotic ghostbursting. The region bounded by PD1 is inherently multistable, and it overlaps with other regions. **(C)** The two-parameter bifurcation of the biophysical model with respect to *I*_*app*_ and the maximum conductance of NMDA receptors (*g*_*NMDA*_), highlighting the various regions of behaviour that the model possesses, including quiescence, tonic firing and doublet/chaotic ghostbursting. **(D)** A sample of three principal firing patterns: tonic spiking at high (1), and low frequencies (2), ghostbursting in the form of doublets (3) and chaotic ghostbursting (4) at different values of *I*_*app*_. These patterns demonstrate the rich repertoire of activity captured by the biophysical model.

At larger *I*_*app*_, the upper unstable branch stabilizes through a supercritical Hopf bifurcation (HB; Fig. 4A) where two envelopes of stable limit cycle emerge (green solid lines). HB divides the parameter space into a tonic firing regime to the left and a second quiescent regime (known as the depolarization block) to the right. In the tonic firing regime, the firing frequency of limit cycles increases with *I*_*app*_. At lower values of *I*_*app*_ near SN1 (Fig. 4A, inset), envelopes of limit cycles undergo a period-doubling (PD1) bifurcation, leading to the formation of the ghostbursting regime to the left of PD1. Further decreasing *I*_*app*_ gives rise initially to a secondary period-doubling (PD2) bifurcation and two saddle-node of periodics bifurcations (labeled SNP1 and SNP2) followed by a cascade of infinite number of bifurcations that become increasingly hard to discern (Fig. 4A, inset); this cascade reflects the onset of doublet firing and chaotic dynamics and the emergence of multistable bursting orbits. Interestingly, to the left of SN1, the model shows bistability, where the quiescent state coexist with bursting orbits, which themselves represent a multistable regime, further highlighting the complex nature of this regime (Supplementary Fig. 4B).

To explore how these distinct dynamic regimes are modulated by intrinsic mechanisms, we next conducted a two-parameter bifurcation analysis in which *I*_*app*_ (vertical axis) was varied in conjunction with the maximal conductances of either *I*_*SK*_: *g*_*SK*_, or *I*_*NMDA*_: *g*_*NMDA*_ (Fig. 4B and C, respectively). This was done by tracking the bifurcation points identified in the one-parameter bifurcation with respect to *I*_*app*_ (Fig. 4A) as a second parameter (*g*_*SK*_, *g*_*NMDA*_) was varied. In the case of the *g*_*SK*_, the continuation in the two parameter space identified four distinct regions of behavior, including no spiking (Fig. 2B; gray, quiescent bottom and depolarization block top), doublet firing (Fig. 2B and D:3; dark green), tonic firing (Fig. 2B and D:1-2; light green), and bursting (Fig. 2B and D:4; purple). Specifically, our results showed that increasing *g*_*SK*_ gradually shifts the boundaries of bursting region towards higher *I*_*app*_ values, indicating that stronger *g*_*SK*_ modulates the spiking activity in the model by reducing the firing rate. Indeed, at intermediate *I*_*app*_values, increasing *g*_*SK*_ can shift the cell from bursting to tonic firing with a lower spiking frequency (Fig. 4D:1-2). On the other hand, larger values of *I*_*app*_ give rise to transition from tonic firing to either doublet firing or bursting, depending on the value of *g*_*SK*_, followed by a return to tonic firing with high frequencies until crossing HB, beyond which the cell ceases to fire at the depolarization block region.

Similarly, the dynamical transitions obtained by continuing the bifurcation points from Figure 4A in a two-parameter space defined by *I*_*app*_ and *g*_*NMDA*_ revealed very rich dynamics (Fig. 4C). As was the case for *g*_*sk*_, several distinct regions of behavior were identified, ranging from quiescence (gray, representing no spiking regions as well as a non-physiological region with unrealistic conductance values), doublet firing (dark green), bursting (purple), and tonic spiking (light green). The bifurcation boundaries, corresponding to SN1, SN2, HB, PD1, SNP1 and SNP2, display a more complex structure compared to the continuation for *g*_*SK*_, reflecting intricate interactions between *I*_*app*_ and *g*_*NMDA*_. Notably, increasing *g*_*NMDA*_ significantly alters the *I*_*app*_ thresholds at which transitions between firing regimes occur. For example, the *I*_*app*_ range that allows for bursting (purple region) is highly sensitive to *g*_*NMDA*_, expanding and shifting as *g*_*NMDA*_ increases. This sensitivity may enable synaptic inputs to more effectively (and robustly) induce bursting. Indeed if *I*_*SK*_ and *I*_*NMDA*_ are blocked (*g*_*SK*_ = 0, *g*_*NMDA*_ = 0), the model exhibits prolonged bursts with prominent depolarizing afterpotentials (DAPs), reverting to dynamics similar to the original *in vitro* ghostbursting model (Supp. Fig. 4C).

Taken together, these results showcase how adjusting *g*_*SK*_ or *g*_*NMDA*_ in conjunction with the applied current governs the transitions among quiescence, tonic firing, and bursting. In particular, our results highlight the crucial role of SK channels and Ca^2+^ dynamics, mediated by NMDA receptors and Ca^2+^ release from the ER, towards shaping firing activity.

### A simplified phenomenological model of burst firing based on a modified Hindmarsh-Rose formalism

Thus far, we have demonstrated that our biophysically detailed model accurately reproduces the action potential shape and the *in vivo* temporal firing patterns of ELL pyramidal cells. While biophysical modeling provides valuable insights into the mechanisms underlying neuronal spiking activity, simpler phenomenological models often offer a better framework for studying circuit-level neural dynamics as well as giving better understanding of which combinations are most relevant (Hindmarsh and Rose, 1984; Laing and Longtin, 2002; Izhikevich, 2003). We developed a new simplified model based on a modified Hindmarsh-Rose (HR) formalism to account for the diverse firing patterns exhibited by these cells (see Methods). Unlike the previously introduced biophysical model, which provides detailed representation of the underlying ionic currents and Ca^2+^ dynamics in full details, the 4-dimensional modified HR model is phenomenological in nature, capable of emulating the key features of ELL pyramidal cell activity, including slow adaptation and intrinsic firing patterns, making it a computationally efficient model to study circuit dynamics of ELL pyramidal cells. By introducing adaptation through a slow variable (*z*) and adaptation current (*u*), the modified HR model accurately replicates gamma-frequency burst oscillations observed in the full biophysical model, as well as the broad range of firing patterns seen in experimental recordings as shown below.

In order to validate the model’s ability to reproduce the diverse firing patterns and chaotic bursting dynamics of ELL pyramidal cells, we performed bifurcation analysis with respect to the applied current (*I*_*app*_) (Fig. 5A). For lower values of *I*_*app*_, the model possesses a branch of stable equilibria (solid orange), representing the resting or holding potential of the cell. As *I*_*app*_ increases, this branch eventually loses stability at a subcritical Hopf bifurcation (HB1), forming a branch of an unstable equilibria (dashed black); this latter branch then folds twice at two saddle-nodes bifurcations (SN1 to the right and SN2 to the left) until it crosses another supercritical Hopf bifurcation (HB2), forming a stable branch of equilibria (solid orange) representing the depolarization block. At HB1, envelopes of unstable periodic orbits (dashed blue) emerge and terminate at a homoclinic bifurcation (HM1; Fig. 5A, top inset). Prior to the SN1, a very small multistable regime of equilibria (solid orange) and periodic orbits (solid green) is also present; this regime is bounded by a supercritical Hopf bifurcation (HB3) to the left, and a subcritical Hopf bifurcation (HB4) to the right; envelopes of small amplitude stable periodic orbits (solid green) emerge from HB3 and eventually undergo period doubling bifurcation (PD1) that gives rise to chaotic small amplitude oscillatory dynamics (dashed green; Fig. 5A, bottom inset) delimited to the left by a cascade of bifurcations that are hard to discern due to their close proximity to each other. Similarly, envelopes of unstable limit cycles (dashed blue) also emerge from HB4 and terminate at a Hopf bifurcation (HB5) to the left of SN1 (Fig. 5A, bottom inset). In a similar fashion, envelopes of unstable limit cycles (dashed blue) also emerge from a subcritical Hopf bifurcation (HB6) to the right of SN2 and terminate at a homoclinic bifurcation (HM2; Fig. 5A). Finally, the envelopes of stable periodic orbits (solid green) that emerge from HB2 merge with envelopes of unstable periodic orbits at a saddle-node of periodics bifurcation (SNP1), which in turn merges with envelopes of stable periodic orbits (solid green) at a saddle-node of periodics bifurcation (SNP2; Fig. 5A). The latter envelopes initially undergo torus bifurcation (TS) to the right and period doubling bifurcation (PD2) to the left, giving rise to chaotic dynamics (Fig. 5B) responsible for the ghostbursting behavior; they eventually terminate to the left by a cascade of infinite bifurcations that become increasingly hard to resolve (Fig. 5A, bottom inset). At PD2, model dynamics transition from tonic firing (Fig. 5C, left) on the right of PD2 to ghostbursting with a varying number of spikes per burst (Fig. 5C, middle and right) on the left, in a manner identical to the biophysical model, allowing for gamma-frequency burst oscillations. The continuation analysis reaching from PD2 into the chaotic bursting regime to the left reveals multistability, eventually terminating into the chaotic attractor regime (rather than a homoclinic bifurcation). Furthermore, for values of *I*_*app*_ below the threshold for spiking originating from HB1, the model exhibits sensitivity to initial conditions, producing either no spiking or single spiking events before returning to the stable resting state reminiscent of type IV excitability (Supplementary Fig. 5).

**Figure 5.**
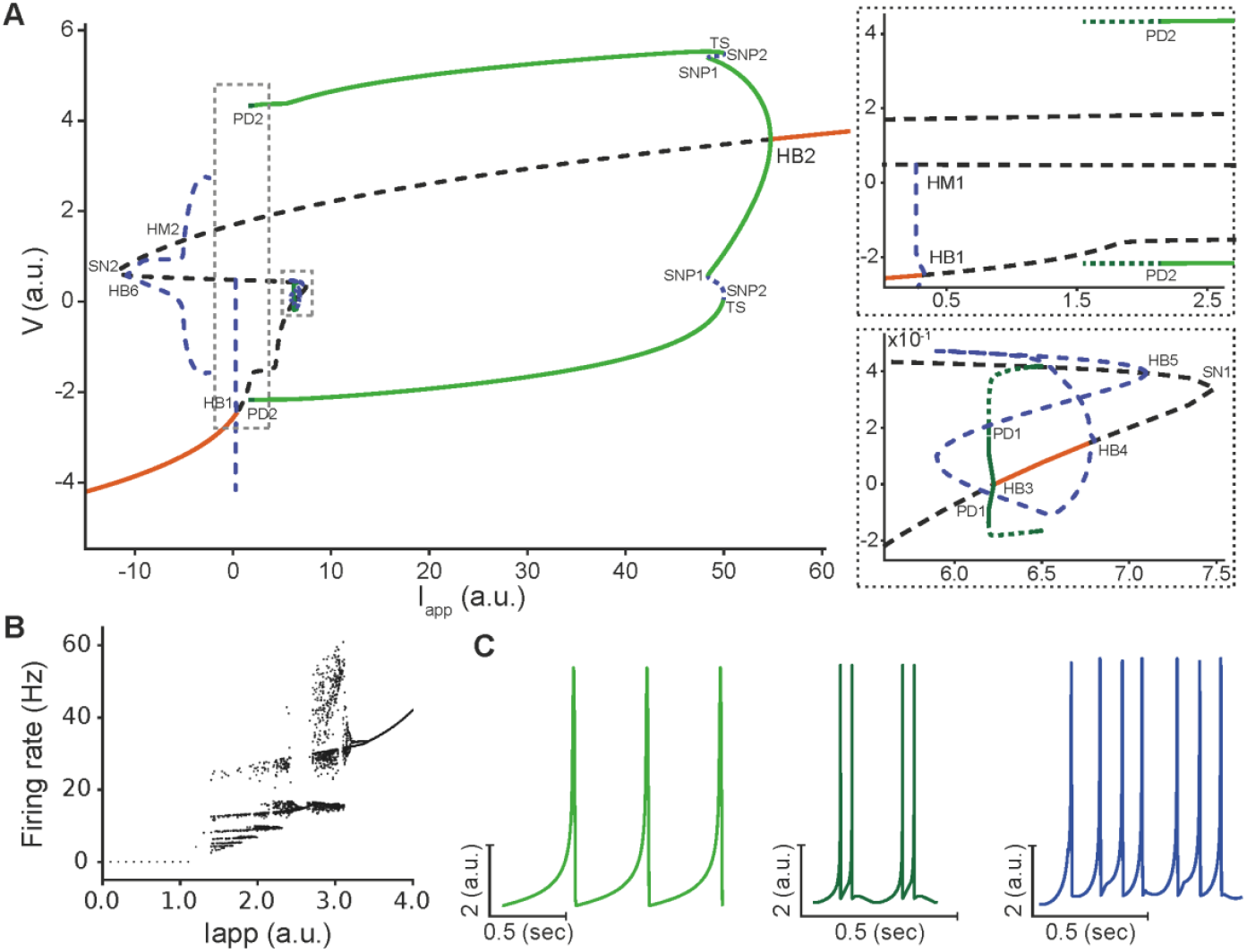
Bifurcation analysis of the deterministic modified Hindmarsh-Rose (HR) model shows similar dynamic regimes as those of the full model. **(A)** One-parameter bifurcation diagram of the voltage variable (*V*) of the modified HR model with respect to the applied current (*I*_*app*_), illustrating the various regimes of behavior corresponding to branches of stable equilibria (orange line), branches of unstable equilibria (dashed black line), envelopes of periodic orbits (green lines) and envelopes of bursting orbits (dashed green line). Within the range of *I*_*app*_ considered, the model undergoes several types of bifurcation points, including 2 saddle-node bifurcations (SN1, SN2), 6 Hopf bifurcations (HB1-HB6), 2 saddle-node of periodics bifurcations (SNP1, SNP2), 2 homoclinic bifurcations (HM1, HM2), 2 period-doubling bifurcations (PD1, PD2) and 1 torus bifurcation (TS), some of which define the boundaries of the various regimes of behavior identified. The two insets provide magnified views of the areas enclosed by the bounding boxes in the main figure, with the top inset corresponding to the larger box and the bottom inset to the smaller box. **(B)** Plot of the firing rate of the modified HR model detected within a time range of 20 seconds long with respect to *I*_*app*_, showcasing the chaotic dynamics exhibited by this model similar to the biophysical model (compare to Supp. Fig. 4B). **(C)** A sample of three principal firing patterns: tonic spiking (*I*_*app*_ = 0.4, left), ghostbursting in the form of doublets (*I*_*app*_ = 1.9, middle) and chaotic ghostbursting (*I*_*app*_ = 1.1, right). These patterns demonstrate the rich repertoire of activity captured by the modified HR model.

Overall, these results illustrate the rich dynamical repertoire of the modified HR model through Hopf, period-doubling, and saddle-node of periodics bifurcations. They highlight the ability of the model to generate ghostbusting reminiscent of gamma-frequency burst oscillations observed in ELL pyramidal cells. One thus would expect that, with noise, the model can transition between stable equilibrium, tonic spiking, and chaotic ghostbursting patterns (Fig. 5B), allowing the model to generate diverse activity patterns that mimic the *in vivo* ELL pyramidal cell activity.

### Stochastic simulations of the modified HR model accurately reproduces ELL pyramidal cell spiking activity seen in vivo

Now that we have demonstrated the ability of the modified HR model to reproduce burst firing dynamics comparable to those observed in the full biophysical model, it remains to check if incorporating stochastic synaptic bombardment can generate the spiking activity of pyramidal cells observed *in vivo*. To do so, we have added extracellular noise term representative of random excitatory and inhibitory inputs to the modified HR model. We then systematically varied its parameters to match the spike train features we obtained from intracellular recordings from ELL pyramidal cells.

Simulating the membrane potential of this example cell model (Fig. 6A, top, teal), alongside the two adaptation variables, *z*(*t*) (middle, red) and *u*(*t*) (bottom, orange) that were introduced to capture the slower timescales of burst onset and termination, showed that, despite the model’s phenomenological nature, it can reproduce the key features of the spiking dynamics of pyramidal cells observed experimentally; that includes the alternation between burst episodes and isolated spiking.

**Figure 6:**
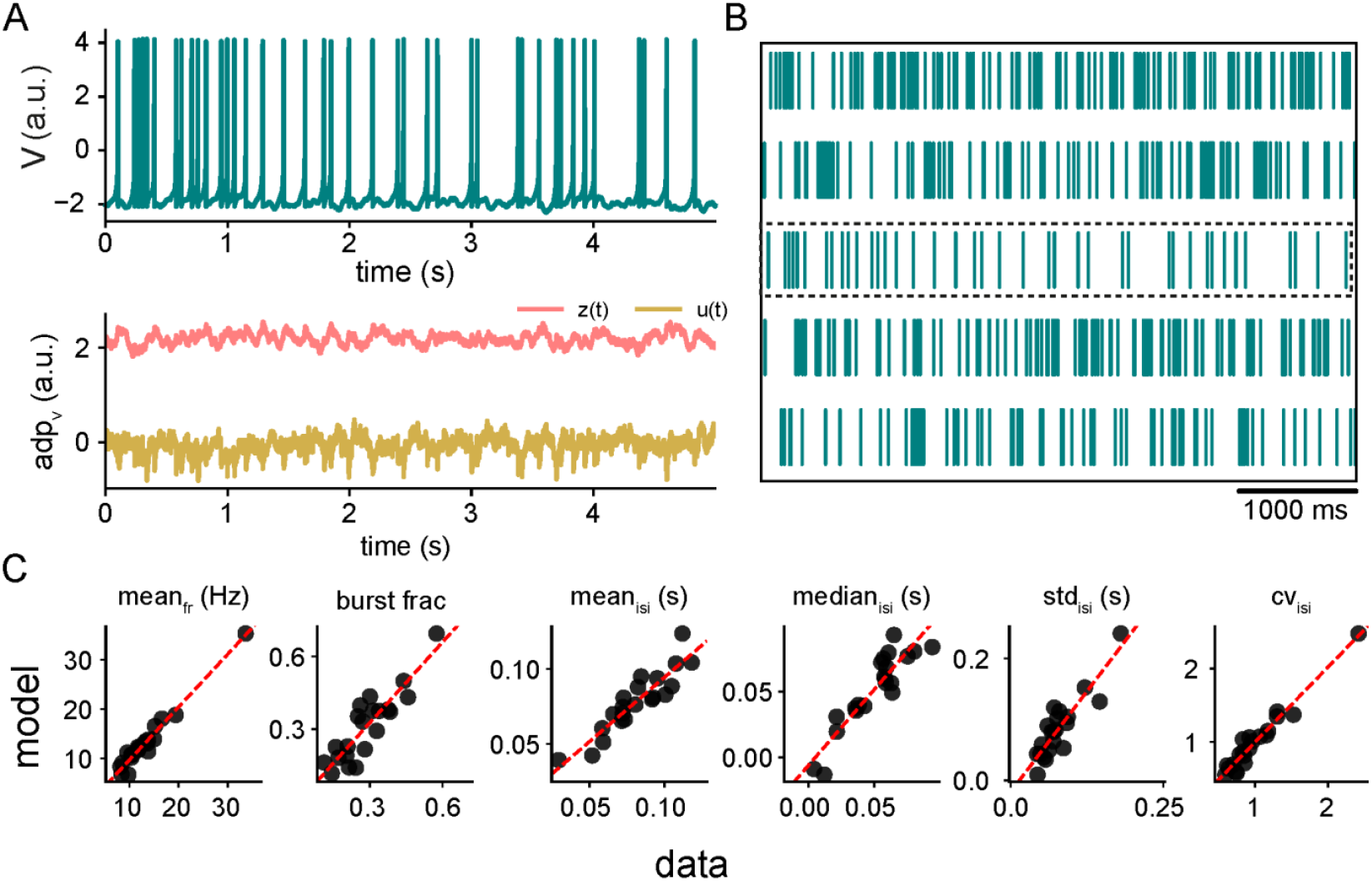
Impact of stochastic synaptic input on the deterministic dynamics of the modified Hindmarsh-Rose (HR) model. **(A)** Simulation of the voltage variable (*V*) of the modified HR model (top, teal), along with the slow adaptation variables *z* (red) and *u* (yellow), for an example ELL pyramidal cell receiving stochastic synaptic input (see Methods). **(B)** Raster spike trains obtained from simulations of the modified HR model for five different representing cells receiving stochastic synaptic input. These spike trains demonstrate the diversity of firing patterns across cells. **(C)** Statistical comparisons between experimental data (dark blue) and model simulations (teal) for spike train features of all recorded ELL pyramidal cells. From left to right: mean firing rate (mean_fr;_ r = 0.99, p=5.07 × 10^−15^), burst fraction (burst frac; r = 0.96, p=8.8 × 10^−8^), mean (mean_isi;_ r = 0.95, p=9.2 × 10^−9^), median (median_isi;_ r = 0.93, p=1.01 × 10^−8^), standard deviation (std_isi;_ r = 0.98, p=6.0 × 10^−8^) and the coefficient of variation (CV_isi_; r = 0.99, p=8.5 × 10^−14^) of ISIs, respectively.

To assess how well the modified HR model captures variability across different cells, we simulated multiple parameter sets corresponding to distinct ELL pyramidal cells. The raster plots of the resulting spike trains for five representative model cells obtained by varying parameter values in the model (Fig. 6B; see Methods) revealed a wide range of firing patterns from predominantly bursting to more isolated spiking, similar to observed *in vivo* activity. We then performed a statistical comparison between the experimental data and model simulations for key spike train metrics, including the mean firing rate (mean_fr_), the burst fraction, and the mean (mean_isi_), median (median_isi_), standard deviation (std_isi_), and coefficient of variations (CV_isi_) of the interspike intervals (Fig. 6C). The scatter plots show a strong correspondence between the model simulations and the experimental data across all metrics (Fig 6C; mean_fr_ r = 0.99, p=5.07 × 10^−15^; burst fraction r = 0.96, p=8.8 × 10^−8^; mean_isi_ r = 0.95, p=9.2 × 10^−9^; median_isi_ r = 0.93, p=1.01 × 10^−8^; std_isi_ r = 0.98, p=6.0 × 10^−8^; and the CV_isi_; r = 0.99, p=8.5 × 10^−14^ of ISIs, respectively), with high correlation coefficients and highly significant p-values. These results indicate that, indicating that despite its phenomenological nature, the model qualitatively reproduces realistic spiking behavior, closely matching both the dynamics of the detailed biophysical model and the experimental intracellular recordings *in vivo*.

In summary, the modified HR model provides a parsimonious yet effective framework for capturing the complex firing behaviors of ELL pyramidal cells *in vivo*. By combining a minimal set of adaptation variables with stochastic synaptic input, the modified HR model reproduces the principal burst dynamics and spiking variability documented experimentally, thereby offering a useful tool for future studies of network-level interactions and higher-order processing in the electrosensory system.

### Parameter sensitivity analysis highlights the importance of adaptation variables and stochastic noise in spiking dynamics

To investigate how the intrinsic parameters of the modified HR model contribute to the observed spiking activity, we conducted a parameter sensitivity analysis using Sobol indices (see Methods). Violin plots were used to visualize parameter distributions, where higher Sobol indices indicate greater influence on model dynamics (Fig. 7A). This analysis identified the top parameters influencing model behavior, with the inhibitory coefficient of slow adaptation variable (*γ*_*d*_) and stochastic noise intensity of the secondary adaptation variable (*σ*_*u*_) showing the highest sensitivity indices. Specifically, we found that *γ*_*d*_ contributes to the transition between tonic and burst firing modes, while *σ*_*u*_ introduces variability that closely aligns the model with *in vivo* spiking patterns. These results suggest that the interplay between intrinsic slow adaptation dynamics and stochastic extrinsic input is critical for reproducing the burst firing patterns and variability seen in experimental data.

**Figure 7:**
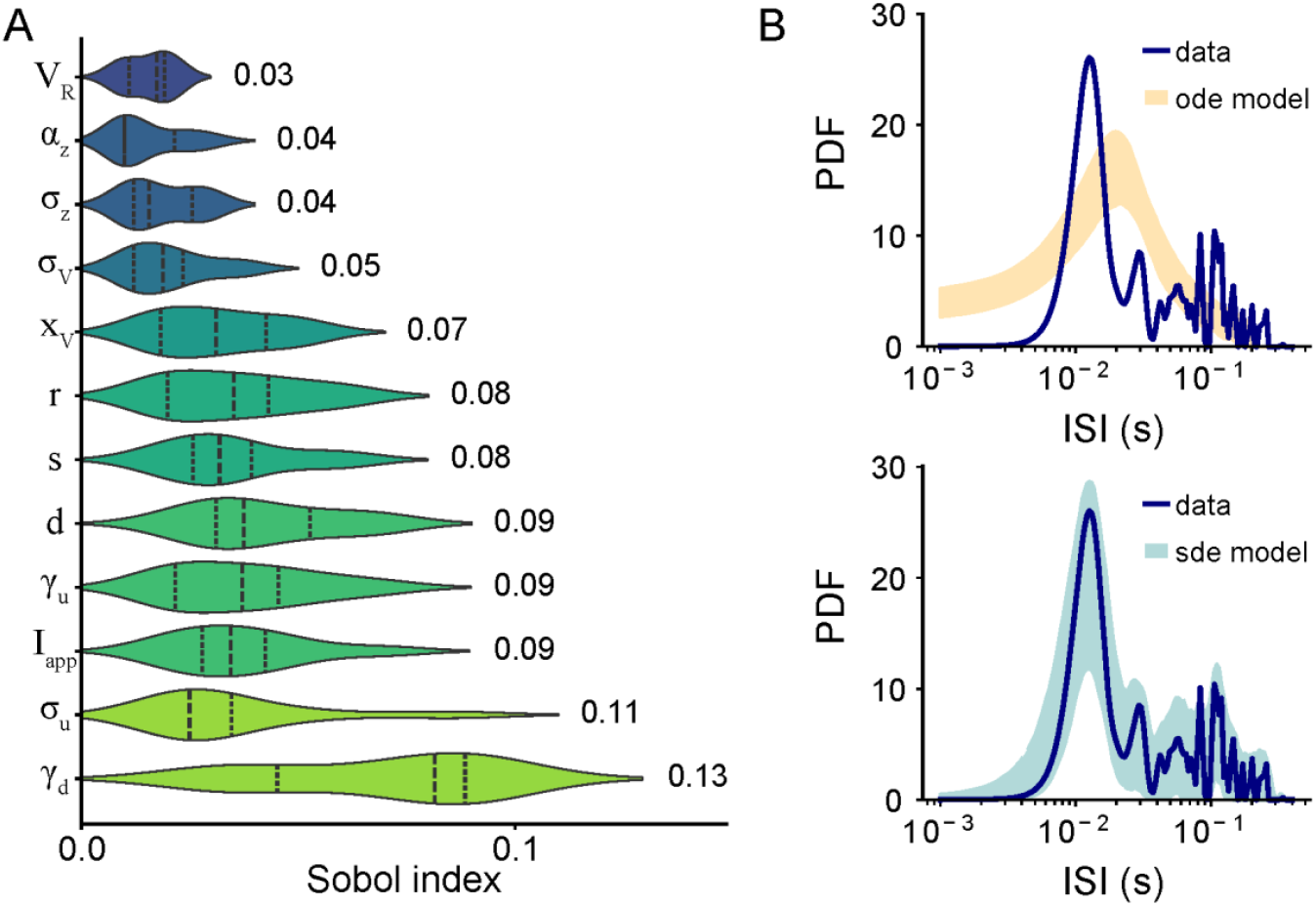
Parameter sensitivity analysis of the modified Hindmarsh-Rose (HR) model reveals significant contribution of the slow adaptation variable and stochastic background noise in aligning it with in vivo spiking activity of ELL pyramidal cells. **(A)** Sobol sensitivity indices for the top 10 model parameters in the modified HR model receiving stochastic synaptic input. The violin plots represent the distributions of parameter values, scaled by their importance as determined by the Sobol sensitivity indices for each parameter. The most sensitive parameters, based on higher Sobol indices, are *γ*_*d*_ and *σ*_*u*_, indicating the greater influence of slow adaptation variable on model behavior compared to other parameters. **(B)** Comparison of the interspike interval (ISI) distributions between an example recorded ELL pyramidal cell (blue) and ISIs from model simulations without (orange, top) and with (teal, bottom) stochastic synaptic input, highlighting the significantly improved alignment of the model with experimental data in the presence of stochastic synaptic input (Kolmogorov-Smirnov test vs data: D_ode model_=0.46, p_ode model_=0.0036; D_sde model_= 0.146 p_sde model_=0.778).

Finally, to assess the role of the deterministic (i.e., intrinsic) components of the model in reproducing the temporal dynamics of spiking activity patterns observed *in vivo*, and their interaction with stochastic synaptic input, we fixed all parameters of the deterministic modified HR model based on the same example ELL pyramidal cell used earlier and simulated its activity with and without stochastic synaptic noise (Fig. 7B). In the absence of stochastic input (top, orange), the model does not capture the burst firing dynamics seen experimentally, as quantified by significant differences between the ISI distributions obtained from the model and from our experimental data (Kolmogorov-Smirnov test vs data: D_ode model_=0.46, p_ode model_=0.0036). However, in the presence of stochastic background input (bottom, teal), the ISI distribution obtained from the model aligns closely with that of the recorded cell (shaded dark blue; Kolmogorov-Smirnov test vs data: D_sde model_= 0.146 p_sde model_=0.778).

To further evaluate the importance of model structure, we simulated the classic HR model using the same parameters as in the modified version. The classic HR model produces an ISI distribution that differs significantly from experimental data (Supp. Fig. 6; Kolmogorov-Smirnov test vs data: D_HR_=0.383, p_HR_= 0.0065). Specifically, it exhibits a rightward shift, fails to capture fast bursting events, and lacks the characteristic bimodality observed *in vivo*. Additionally, the distribution is noticeably narrower, indicating reduced variability. These results highlight that the modifications introduced in our model, beyond just parameter tuning, are essential for accurately reproducing the complex temporal structure of ELL pyramidal cell spiking. These findings underscore the critical role of stochastic components in modeling synaptic bombardment and the high-conductance states characteristic of *in vivo* conditions.

## Discussion

### Summary of results

Neurons *in vivo* operate under different conditions than *in vitro* as they are constantly bombarded by synaptic inputs as well as neuromodulators that are typically absent *in vitro* (Wei et al., 2023; Koch et al., 2025; Rabuffo et al., 2025). Understanding how these factors interact with the intrinsic neuronal mechanisms is crucial for uncovering the critical computations performed by neural circuits in the behaving animal. In the present study, we addressed this gap by investigating how *in vivo* conditions affect the intrinsic burst firing displayed by ELL pyramidal cells *in vitro*. To do so, we developed a two-compartment biophysical model that combines the membrane voltage dynamics with Ca^2+^ mobilization. After fitting the model parameters against intracellular recordings, we first demonstrated that the model successfully captures the intrinsic and extrinsic mechanisms underlying the firing patterns of these cells. Next, we used bifurcation analysis to identify how specific conductances, such as SK (*g*_*SK*_) and NMDA (*g*_*NMDA*_), shape transitions between quiescent, tonic, and bursting regimes, revealing complex multistable dynamics. We then designed a simplified model based on the modified Hindmarsh-Rose (HR) formalism to phenomenologically reproduce the firing patterns of ELL pyramidal cells seen *in vivo*. This model retained key dynamical features, including chaotic bursting, while offering computational efficiency, providing a robust framework for exploring spiking activity of these cells. Finally, the parameter sensitivity analysis highlighted the critical role of slow adaptation mechanisms and stochastic input in shaping spiking patterns detected *in vivo*. The two models together offer a mechanistically grounded framework for understanding how intrinsic dynamics and *in vivo* conditions interact to shape the activity of ELL pyramidal cells.

### The Impact of dendritic integration and stochastic synaptic input on burst firing

Seminal *in vitro* studies of ELL pyramidal cells described a “ping-pong” mechanism driven by the backpropagation of somatic action potentials into the dendrites This process leads to dendritic spikes and subsequent somatic depolarizing afterpotentials (DAPs) that trigger further spikes until they eventually terminate due to dendritic failure (Lemon and Turner, 2000; Doiron et al., 2001; Rashid et al., 2001; Doiron et al., 2002). In contrast, *in vivo* recordings, including those presented here, consistently show bursts that are typically shorter, often lack prominent DAPs or pronounced afterhyperpolarizations (AHPs), and exhibit more variable interspike intervals (ISIs) (Bastian and Nguyenkim, 2001; Oswald et al., 2004; Toporikova and Chacron, 2009; Metzen et al., 2016). Our biophysical modeling provides a novel mechanistic insight to explain these discrepancies by incorporating key features unique to *in vivo* conditions. Specifically, the model demonstrates that dendritic Ca^2+^ influx, primarily mediated by NMDA receptor dynamics, activates SK channels whose hyperpolarizing effects act to prematurely terminate bursts before the full development of the ping-pong mechanism (Toporikova and Chacron, 2009). This truncation shortens burst duration and reduces DAP prominence, resulting in burst dynamics that closely align with those observed *in vivo*. Indeed, this result is consistent with experimental studies showing that pharmacological inhibition of SK channels by apamin (Ellis et al., 2007b), or blockade of intracellular Ca^2+^ using BAPTA (Krahe et al., 2008; Toporikova and Chacron, 2009), restore long-duration bursts with prominent DAPs *in vivo* by preventing the premature termination of bursts. Complementing this gating mechanism is the influence of extrinsic synaptic input, which introduces continuous membrane potential variability. This variability contributes to irregular burst timing and disrupts the structured burst patterns observed *in vitro*, which otherwise exhibit strong correlations with specific stimulus properties (Destexhe et al., 2003; Avila-Akerberg and Chacron, 2011; Hua et al., 2025). By applying parameter sensitivity analysis, we further highlighted in this study the importance of these factors by revealing that both slow adaptation dynamics and stochastic noise intensity are dominant determinants of the observed variability and structure of *in vivo* burst events, suggesting a crucial interplay between intrinsic cellular properties and the presence of stochasticity caused by active synaptic bombardment. This dynamic interplay plays a key role distinguishing *in vivo* burst generation from those firing patterns seen *in vitro*.

### Modulation of firing dynamics by feedback and neuromodulator serotonin

It is well-known that ELL pyramidal cells *in vivo* are subject to substantial neuromodulatory (Deemyad et al., 2013; Marquez and Chacron, 2018; Marquez and Chacron, 2023) and synaptic input (Bastian and Nguyenkim, 2001; Huang et al., 2019), which can dynamically influence their spiking activity and burst firing patterns over time. Within the ELL, neuromodulatory inputs, including acetylcholine (Ellis et al., 2007a; Mehaffey et al., 2008a), and serotonin (Larson et al., 2014; Marquez and Chacron, 2020b, a) are known to broadly regulate neuronal function by modifying excitability and responsiveness to stimulation. Our models provide a robust framework for understanding how such factors modulate firing dynamics in ELL pyramidal cells by exploiting the cell’s intrinsic multistability. For instance, it has been shown experimentally that serotonin released from raphe nuclei inhibits SK channels, reducing the medium afterhyperpolarization and promoting burst firing in ELL pyramidal cells (Deemyad et al., 2011). SK channels have also been shown to play a critical role in regulating neuronal excitability and frequency tuning (Ellis et al., 2007b). Our model builds on these findings by quantitatively predicting that reducing SK conductance (*g*_*SK*_) lowers the burst threshold and expands the bursting regime (Deemyad et al., 2013; Metzen et al., 2016; Marquez and Chacron, 2020a; Akhshi et al., 2023). The model can further be used to explore how serotonin affects membrane excitability and promotes burst firing, as seen experimentally, by leveraging the intrinsic dynamics (e.g., SK channels), thereby offering mechanistic insight into experimental results showing serotonin modulation enhances the encoding of behaviorally relevant stimuli such as specific communication signals (Deemyad et al., 2013; Marquez and Chacron, 2018).

In addition to neuromodulation, *in vivo* spiking activity of neurons is significantly affected by feedback from higher brain areas, which can reflect behavioral state, attention, or predictive processing (Briggs et al., 2013; Zagha and McCormick, 2014). Such feedback can dynamically regulate ELL pyramidal cell excitability and responsiveness to sensory input (Lemon and Turner, 2000; Bastian and Nguyenkim, 2001; Toporikova and Chacron, 2009) through both stochastic synaptic bombardment (Avila-Akerberg and Chacron, 2011) and NMDA receptor-mediated modulation of membrane potentials (Bastian, 1993; Berman et al., 2001; Mease et al., 2014). Our bifurcation analysis demonstrates that changes in NMDA conductance (*g*_*NMDA*_) strongly influences the boundaries between firing regimes, thereby offering a dynamical basis for the established ability of feedback pathways to control whether a neuron operates in silent, tonic, or bursting modes (Bastian and Nguyenkim, 2001). Even brief or subtle shifts in feedback or neuromodulatory factors, according to our model, can shift cells across these regime boundaries, consistent with experimental observations (Deemyad et al., 2013; Marquez and Chacron, 2018, 2020a). Furthermore, the continuous presence of stochastic synaptic activity *in vivo* introduces additional membrane potential variability and can enhance the responsiveness of pyramidal cells to changes in feedback and neuromodulation (Destexhe et al., 2003; Prescott et al., 2008; Avila-Akerberg and Chacron, 2011). Together, these mechanisms provide a flexible means by which the firing mode of ELL pyramidal cells can be rapidly and reversibly modulated via neuromodulation (Ellis et al., 2007a; Mehaffey et al., 2008b; Deemyad et al., 2013; Marquez and Chacron, 2020a), feedback strength (Bastian and Nguyenkim, 2001; Berman et al., 2001; Metzen and Chacron, 2023), and ongoing synaptic activity (Avila-Akerberg and Chacron, 2011; Huang et al., 2019). Our model thus provides a biophysical basis for the observed dynamic control of burst firing *in vivo* and explains how physiological modulation can drive transitions between firing states.

### Multistability: A mechanism for adaptive neuronal computation

A key finding from our bifurcation analysis is the identification of complex dynamical behaviors characterized by multistability in both models. The biophysical model exhibits multistability near the saddle-node (SN1) and period doubling (PD1) bifurcations, where quiescent states can coexist with bursting activity, reflecting a robust capacity for state transitions under the same depolarizing current (Malashchenko et al., 2011). Furthermore, the chaotic bursting regime itself exhibits multiple coexisting bursting patterns (multistability within bursting), reflecting the complex attractor structures that are characteristic of neurons with slow Ca^2+^-related dynamics (Malashchenko et al., 2011; Iosub et al., 2015). Similarly, the modified HR model displays multistable bursting dynamics (near PD2), further illustrating the model’s capability to reproduce the complex firing dynamics observed in the biophysical model. This multistability has significant implications for neuronal coding (Bertram et al., 1995; Newman and Butera, 2010; Buchin et al., 2016), facilitating for example rapid and context-dependent switching between different firing modes, such as quiescent, tonic spiking, and bursting, without requiring long-term synaptic changes. Despite the stochastic nature of synaptic inputs, the intrinsic properties of dendritic and axonal compartments enable neurons to generate and differentiate between single spikes and bursts, assigning distinct coding roles to each firing mode (Spruston, 2008; Larkum et al., 2009; Major et al., 2013). While these coding roles arise from the compartmental integration and active properties of neurons, synaptic noise can act as a perturbation that pushes the neurons across firing regime boundaries within its multistable landscape, even though it doesn’t directly encode stimulus features (Doiron et al., 2001; Doiron et al., 2003; Akerberg and Chacron, 2011; Akhshi et al., 2023). For instance, in the subthreshold stable node regime of both models, a transient perturbation can elicit a single spike consistent with Type IV excitability (Mitry et al., 2020; MacKay et al., 2021). The mechanism typically involves the neuron being poised near a bifurcation (such as a SNIC or a homoclinic bifurcation), where a brief input allows a single spike excursion before returning to the stable resting state. Such single-spike excitability can be leveraged as a precise temporal coding mechanism, enabling neurons to transmit information with high temporal fidelity by signaling the exact onset of a stimulus or encoding fine temporal features through isolated spikes (Mitry et al., 2020). Indeed, this capability arises from the rapid and temporally precise nature of single spikes, allowing them to encode fine temporal patterns that are critical for sensory discrimination (Mainen and Sejnowski, 1995; Oswald et al., 2004; Metzen et al., 2025). In contrast, bursts offer a robust means of signaling in noisy environments and may encode not only more generalized or invariant stimulus attributes (Krahe and Gabbiani, 2004; Oswald et al., 2007; Avila-Akerberg and Chacron, 2011; Metzen et al., 2016), but also different time scales (Bertram et al., 1995).

Taken together, this complex dynamical landscape, marked by the coexistence of multiple firing states, can enable neurons to dynamically adjust their firing mode in response to transient inputs or ongoing synaptic fluctuations. Such multistability thus serves as a powerful mechanism for adaptive computation, allowing neurons to flexibly alter their coding strategies in accordance with changing sensory contexts or behavioral demands (Gjorgjieva et al., 2016).

### Coding by burst firing in vivo

The functional significance of burst firing in ELL pyramidal cells *in vivo* diverges markedly from classical interpretations derived from *in vitro* experiments. *In vitro*, bursts have been associated with the precise encoding of stimulus features, such as slope or intensity, via strong correlations between burst structure and specific stimulus characteristics (Lemon and Turner, 2000; Doiron et al., 2001; Kepecs et al., 2002; Oswald et al., 2007). However, pervasive synaptic bombardment *in vivo* introduces both a high conducting state in neurons (Koch et al., 2025) and a substantial spike timing jitter, which can obscure or degrade the information conveyed by burst patterns and reduce their reliability for downstream decoding (Destexhe et al., 2003; Fellous et al., 2003; Akerberg and Chacron, 2011). Recent *in vivo* studies indicate that ELL bursts act as robust and flexible event markers (Akerberg and Chacron, 2011; Deemyad et al., 2013; Metzen et al., 2016; Metzen et al., 2025). They are particularly effective for signaling the occurrence of low-frequency, behaviorally relevant stimuli rather than encoding the exact details of stimulus waveforms (Oswald et al., 2004; Metzen et al., 2016; Huang et al., 2019; Wang and Chacron, 2021; Metzen et al., 2025). The sparsity and distinctiveness of bursts, as opposed to ongoing single spikes, makes them well-suited to flag salient events in complex sensory environments as burst events are often more resilient to the noise inherent in the *in vivo* state (Krahe and Gabbiani, 2004; Shao et al., 2021). Additionally, bursts in ELL pyramidal cells contribute to adaptive computations beyond sensory detection. They play a direct role in regulating feedback plasticity, helping the system maintain appropriate gain control and selectivity for external stimuli (Harvey-Girard et al., 2010; Bol et al., 2011). As mentioned earlier, the propensity for burst firing is also subject to modulation by neurotransmitters like serotonin, which can shift the balance of excitability and enhance responses to social signals by suppressing SK channel activity (Deemyad et al., 2013). On a network level, correlated burst firing supports synchronous activity across populations, which may facilitate more reliable population coding and enhance the robustness of signal detection (Chacron and Bastian, 2008; Simmonds and Chacron, 2015; Metzen and Chacron, 2023; Metzen et al., 2025). Thus, the *in vivo* function of bursts is not primarily to encode fine stimulus structure, but to provide a flexible and resilient mechanism for highlighting salient, behaviorally relevant events, supporting plasticity, and enabling robust communication in a noisy and dynamically changing environment.

### Generalizability and implications for other systems

The dual-modeling approach employed in this study, using a detailed biophysical model for mechanistic grounding alongside a simplified phenomenological model for analytical tractability and computational efficiency, can be applied to other cell types to bridge the gap between *in vitro* and *in vivo* dynamics (Destexhe et al., 2003; Prescott et al., 2008; Prescott and Sejnowski, 2008; Wei et al., 2023; Koch et al., 2025). This integrated framework allows direct testing of specific ionic mechanisms (e.g., SK channel roles), while facilitating systematic analysis (e.g., bifurcation) and large-scale simulations via the efficient modified HR model. The mechanisms identified here for *in vivo* spiking activity and burst dynamics likely extend beyond the electrosensory system. While burst firing is a widespread phenomenon across various brain regions (Krahe and Gabbiani, 2004; Shao et al., 2021), the electrosensory system shares notable similarities with other sensory modalities, including the mammalian vestibular (Mackrous et al., 2020; Carriot et al., 2022), visual (Clarke et al., 2015), and auditory (Metzen et al., 2015) systems. Understanding how specific *in vivo* mechanisms, such as the Ca^2+^-related dynamic interactions identified in this study, shape the structure and function of bursts is therefore broadly relevant. Moreover, the multistability uncovered by our models suggests a general computational principle: that dynamic switching between distinct firing modes may support adaptive coding across sensory, motor, and cognitive domains (Drion et al., 2015; Gjorgjieva et al., 2016). Finally, our modeling framework also enables broader applications, such as probing cellular heterogeneity (Nandi et al., 2022; Akhshi et al., 2023), incorporating activity-dependent plasticity (Huang et al., 2019; Metzen and Chacron, 2023), and leveraging the computational efficiency of the modified HR model for large-scale simulations (Litwin-Kumar and Doiron, 2012; Perez-Nieves et al., 2021).

## Materials and Methods

### Experimental setup

The wave-type weakly electric fish *Apteronotus leptorhynchus* (*N = 3*) of either sex was used exclusively in this study. The fish were sourced from commercial tropical fish suppliers and housed in groups ranging from 2 to 10 individuals. Water conditions were carefully maintained at temperatures between 26 and 29°C, with conductivities ranging from 300 to 800 µS·cm^−1^, following established care guidelines (Hitschfeld et al., 2009). All procedures involving animals were reviewed and approved by the McGill University Animal Care Committee (#5285) and were conducted in accordance with the regulations set by the Canadian Council on Animal Care.

### Surgery

Details of the surgical procedures have been previously described in detail (Chacron et al., 2003; Toporikova and Chacron, 2009; Metzen et al., 2015). Briefly, animals were immobilized for electrophysiological recordings via intramuscular administration of tubocurarine (0.1–0.5 mg, Sigma). Fish were then placed in an experimental tank (30 × 30 × 10 cm^3^) filled with water from their home tank and provided with a constant flow of oxygenated water through the mouth at 10 mL/min rate. Subsequently, the animal’s head was locally anesthetized with lidocaine ointment (5%; AstraZeneca, Mississauga, ON, Canada), the skull was partly exposed, and a small window was opened over the hindbrain.

### Recordings

Intracellular recordings were conducted *in vivo* from ELL pyramidal cells of either cell type (ON- or OFF-) using sharp electrodes, following standard procedures (Bastian et al., 2002; Toporikova and Chacron, 2009; Deemyad et al., 2013). Recordings were made under baseline conditions (i.e., in the presence of the animal’s unmodulated EOD). Micropipettes had resistances ranging from 20 to 80 MΩ and were filled with 3 M potassium chloride (KCl), as done previously (Bastian et al., 1993). Pyramidal cells were recorded at depths ranging from 300 to 800 µm below the brain surface, corresponding to the centrolateral and lateral ELL segments based on established anatomical landmarks (Maler et al., 1991; Maler, 2009). Recordings were digitized at 10 kHz using CED 1401 plus hardware and Spike2 software (Cambridge Electronic Design, Cambridge, UK) and stored on a hard drive for further analysis. Overall, we recorded from a total of *n = 20* ELL pyramidal cells across *N=3* fish.

### Computational Model

#### Biophysical model of ELL pyramidal cells

We developed a two-compartmental biophysical model of ELL pyramidal cell activity (Doiron et al., 2002; Toporikova and Chacron, 2009; Akhshi et al., 2023). The model consists of two compartments and is based on the Hodgkin-Huxley formalism. The equations describing the membrane voltages in the somatic (*V*_*S*_) and dendritic (*V*_*D*_) compartments are given by

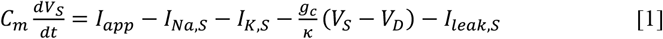

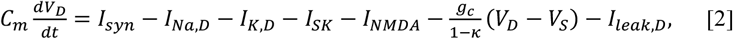

where *C*_*m*_ is the membrane capacitance, *I*_*Na,i*_ and *I*_*K,i*_ are the fast inward Na^+^ and the slow outward delayed rectifier K^+^ currents in the soma (*i* = *S*) and the dendrite (*i* = *D*), respectively, *I*_*app*_ is the depolarizing current to the somatic compartment, *I*_*syn*_ is the synaptic current to the dendritic compartment, *I*_*SK*_ and *I*_*NMDA*_ are the outward Ca^2+^-dependent small-conductance K^+^ and the inward NMDA Ca^2+^ currents acting on the dendrite, respectively, and *I*_*leak*_ is the passive leak current present in both compartments. The two compartments are linked together through a resistor, where *g*_*c*_ is the maximum conductance and *κ* is the somatic-to-dendritic area ratio. The currents *I*_*Na,i*_ and *I*_*K,i*_ (*i* = *S, D*) are necessary to generate somatic action potentials and the proper spike backpropagation that yields somatic depolarizing afterpotentials (DAPs) (Doiron et al., 2002). The two currents *I*_*NMDA*_ and *I*_*SK*_ were added to the dendritic compartment because of the important role they play in regulating spiking activities of ELL pyramidal cells both *in vitro* (Ellis et al., 2007b) and *in vivo* (Toporikova and Chacron, 2009; Huang et al., 2016; Huang and Chacron, 2017). The kinetics of the ionic currents included in each compartment (*i* = *S, D*) are given by

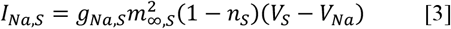

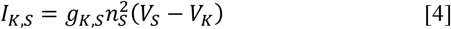

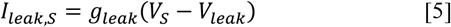

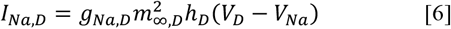

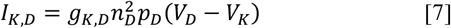

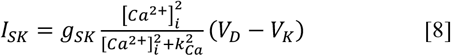

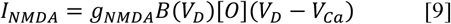

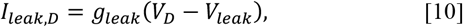

where *g*_*j,i*_ *(j* = *Na, K, leak, SK, NMDA* and *i* = *S, D*) are the maximum conductances, *V*_*j*_ (*j* = *Na, K, Ca, leak*) are the Nernst potentials, [*Ca*^2+^]_*i*_ is the Ca^2+^ concentration in the dendritic compartment (μM), *k*_*Ca*_ is the half-maximum activation of the SK channel (μM), *B*(*V*_*D*_) is the magnesium block, given by (Destexhe et al., 1994)

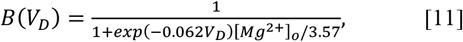

[*Mg*^*2+*^]_*o*_ is the extracellular magnesium concentration (mM), [*O*] is the open probability of the NMDA receptors defined by a Markov model comprised of three closed, one open and one desensitized states with transition rates *R*_*i*_ (*i* = *b, u, c, o, d, r*) between them, *m*_∞,*i*_ *(i* = *S, D)* is the steady state activation of *I*_*Na,i*_, and *x (x* = *n*_*i*_, *h*_*D*_, *p*_*D*_; *i* = *S, D)* are the activation/inactivation gating variables whose steady states and time constants are denoted by *x*_∞_ and *τ*_*x*_, respectively, and whose dynamics are governed by the equation

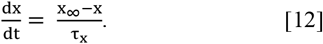

The steady state activation/inactivation functions are given by

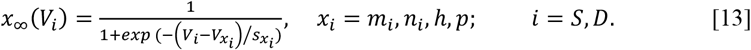

In this model formalism, it was assumed that *I*_*SK*_ and *I*_*NMDA*_ are co-localized within spines. We therefore did not consider Ca^2+^ diffusion within/between spines.

The NMDA receptor activation depends on the concentration of glutamate following principles of ligand-gated channels described previously (Destexhe et al., 1994), where the transition rates between unbound and bound states is based on the concentration of ligand. In our model, the timing of the release of glutamate follows a Poisson process with firing rate *λ*_*glu*_. The glutamate concentration following each release event was described by an alpha function (Destexhe et al., 1994), given by

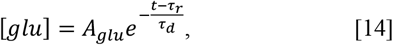

where *τ*_*r*_ is the timing of glutamate release, determined by presynaptic spike times, and *τ*_*d*_ is the time constant of the alpha function, and *A*_*glu*_ is a scaling constant. The parameter *τ*_*d*_ was chosen to be fast enough to prevent glutamate release events from overlapping.

The Ca^2+^ mobilization model followed the flux-balance formalism; it describes fluxes across the cell and ER membranes in the dendritic compartment. Ca^2+^ mobilization across the cell membrane includes three fluxes through the NMDA receptors: *J*_*NMDA*_ = *αI*_*NMDA*_, where 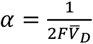(*F* is Faraday’s constant and 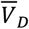 is the volume of the dendritic compartment), plasma membrane Ca^2+^ ATPases (PMCA) pumps: *J*_*PMCA*_, and leak: *J*_*INLeak*_. Ca^2+^ mobilization across the ER membrane, on the other hand, includes three fluxes through the IP3Rs: J_IP3R_, sarco/endoplasmic reticulum ATPases (SERCA) pumps: J_SERCA_, and leak: *J*_*ERLeak*_. The Li-Rinzel model was adopted to describe IP3R kinetics (Li and Rinzel, 1994; Mikolajewicz et al., 2021; Oprea et al., 2022). It is given by

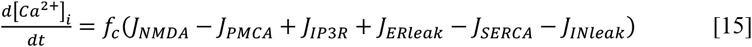

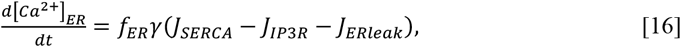

where *f*_*c*_ (*f*_*ER*_) is the fraction of free Ca^2+^ concentration in the cytoslic (ER) component of the dendrite, and γ is the volume ratio of cytosol to ER in the dendrite. The fluxes through the PMCA and SERCA pumps are given by

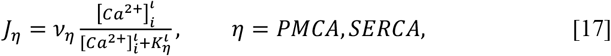

where *ι* = 2 is the Hill coefficient, *ν*_*η*_ are the maximum flux rates (μM/s), and *K*_*η*_ are the half-maximum activations for Ca^2+^ flux (μM). Fluxes due to leak across cell and ER membranes are given by

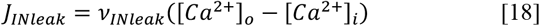

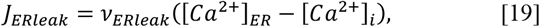

where *ν*_*ξ*_ (*ξ* = *INleak, ERleak*) are the maximum flux rates (μM/s). Finally, flux through IP3Rs is adopted from the Li-Rinzel model (Li and Rinzel, 1994) and is given by

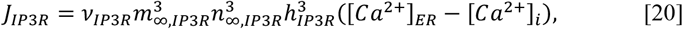

where ν_*IP*3*R*_ is the maximum flux rate (μM/s),

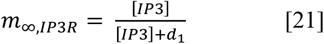

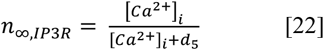

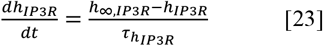

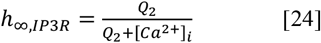

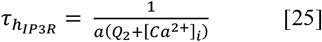

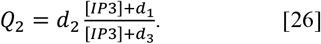

Note that [*IP*3] is the cytosolic concentration of IP3 in the dendrite (μM).

The synaptic input current, *I*_*syn*_, was used to represent all the stochastic background excitatory and inhibitory pre-synaptic activity, defined by *W*_*x*_(*t*) (*x* = *E, I*), applied to the dendritic compartment (Manwani and Koch, 1999a; Brake et al., 2024). It has been previously suggested that the power spectra of excitatory and inhibitory background activity in synaptic inputs exhibit a power law spectrum, *S*_*x*_(*f*)∼ 1/*f*^*β*^, where *β* determines the exponent of steepness of the slope of the power spectrum (Brake et al., 2024; Brake and Khadra, 2025). Based on this, the total synaptic input incorporated into the model can be described by

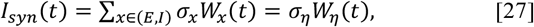

where *σ*_*η*_ is the total noise intensity obtained by integrating the two components, *W*_*η*_(*t*) is the sum of both excitatory and inhibitory pre-synaptic inputs reconstructed using the following expression

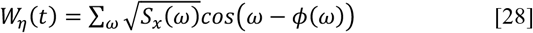

with *ω* = 2*πf* and *ϕ*(*ω*) ∈ [0,2*π*] is a random phase multiplied by the spectrum in the frequency domain. Once the spectrum is reconstructed, the synaptic time series can be obtained using inverse Fourier transform of the spectrum as described previously (Timmer and Koenig, 1995).

#### Modified Hindmarsh-Rose model

The classical Hindmarsh-Rose (HR) model is a well-known three-dimensional system used to study neuronal spiking and bursting behaviors (Hindmarsh and Rose, 1984; Izhikevich et al., 2003). However, in its classical form, this model shares a common limitation with many other phenomenological bursting models in that the intraburst spike frequency typically decreases over the course of the burst cycle. In contrast, ELL pyramidal cells exhibit an increasing spike frequency during the intraburst interval. Furthermore, the classical HR model displays abrupt transitioning from a quiescent state to burst firing as the depolarization current is increased, whereas ELL pyramidal cells exhibit gradual transitions from quiescence to tonic firing before reaching the bursting state in response to increasing current. To address these discrepancies and better capture experimentally observed spiking and bursting patterns in ELL pyramidal cells *in vivo*, we developed a simplified phenomenological model based on the Hindmarsh–Rose (HR) formalism. The classical HR model comprises a single fast variable (*V*) that mimics the membrane potential traces of neurons, a recovery variable (*y*) that controls the rapid recovery processes following action potentials, and a slow adaptation variable (*z*) that reflects the slow adaptation mechanisms that modulate neuronal excitability over longer timescales, regulating the spike frequency. We extended the HR model by introducing a new secondary slow variable (*u*) that modulates the spiking activity through an activation gating mechanism, enabling dynamic regulation of neuronal excitability based on membrane potential. The model is governed by the following set of differential equations:

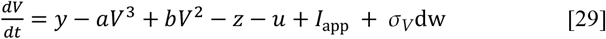

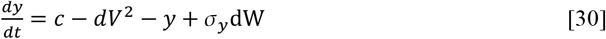

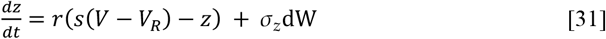

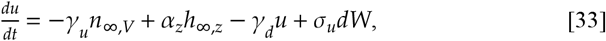

where the dynamics of the newly introduced adaptation variable *u* is governed by the interaction between voltage-dependent activation (*n*_∞,*V*_) and a secondary inactivation process (*h*_∞,*z*_). Moreover, *I*_app_ is an external input current representing the stimulus input to the cell, and noise terms *σ*_*i*_ (*i* = *V, y, z, u*) incorporate all stochastic fluctuations at different time accounting for both intrinsic and external background noise in each equation (Manwani and Koch, 1999b, a).

In this formalism, *n*_∞,*V*_ represents the steady-state activation function governed by membrane potential *V*, where

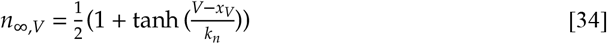

and *h*_∞,*z*_ represents the steady-state inactivation function dependent on the slow adaptation variable *z*,

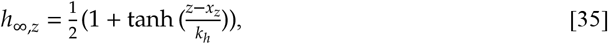

where *x*_*V*_, and *x*_*z*_ denote the baseline values, and *k*_*n*_, *k*_*h*_ represent the slopes of these gating variables, determining how rapidly they respond to changes in *u*, and *z*, respectively. Additionally, dW_*i*_ (*i* = *V, y, z, u*) are independent Wiener processes with zero mean and a standard deviation of 1.

This formulation allows the model to capture adaptive modulation of neuronal excitability, incorporating a dynamic activation-inactivation mechanism that extends the classical HR model. The inclusion of *u* introduces additional complexity that enhances the model’s ability to reproduce experimentally observed spiking and bursting patterns in ELL pyramidal cells.

### Data Analysis

#### Spike detection and feature extraction

Both the data obtained from intracellular recordings and the model simulations were analyzed using a custom code built upon the Intrinsic Physiology Feature Extractor (IPFX) library (https://ipfx.readthedocs.io/) from Allen Brain Institute. Spike detection was performed using a threshold-based algorithm (−30 mV relative to baseline for the data and the ELL pyramidal cell model, and 0 mV for the modified HR model). To avoid artifacts, the maximum allowable time between consecutive spikes was set to 1 ms. Individual action potentials were first isolated from voltage recordings by extracting segments of the membrane potential trace within a ±4 ms window around each spike peak. For each action potential, six AP features were extracted, including voltage threshold for spiking, peak amplitude (peak-to-trough voltage), trough amplitude, voltage at the midpoint of the upstroke phase, voltage at the midpoint of the downstroke phase (voltage at 50% amplitude of the upstroke and downstroke phases), afterdepolarization potential (ADP), and trough amplitude. These features together were used for comparison between the ELL pyramidal cell model and experimental data during optimization. To reduce variability across spikes, the average of each action potential feature across multiple spikes was computed and used for subsequent analyses.

#### Inter-spike interval (ISI) distribution estimates

To estimate ISI distributions, spike times were extracted following spike detection, based on the timing of detected peak amplitudes from either recordings or simulations (5 sec duration). ISIs were calculated as the differences between consecutive spike times. For visualization, histograms of ISI distributions were constructed using 2 ms bin widths. Additionally, kernel density estimation (KDE) was applied to generate a smoothed representation of the ISI distributions, using a Gaussian kernel with bandwidth selected according to Scott’s rule (Scott, 1979). To obtain error intervals around the ISI distributions, we used a bootstrapping approach with 50 resamples, where 4 second segments of the spike time data were randomly subsampled. The variance around the mean KDE estimate across these samples was then calculated and plotted as the error interval.

#### Burst identification and quantification

Burst dynamics were analyzed using custom scripts following established methods in prior studies (Oswald et al., 2004; Avila-Akerberg et al., 2010; Khosravi-Hashemi et al., 2011; Marquez and Chacron, 2023; Metzen et al., 2024). Bursts were identified by setting an ISI threshold of 12 ms, a commonly used value to distinguish intra-burst spikes from isolated spikes in ELL pyramidal cells (Marquez and Chacron, 2018; Marquez and Chacron, 2023). Burst fraction was calculated as the ratio of spikes within bursts to the total spike count. ISI distributions were computed to quantify spiking variability and to characterize the firing patterns as bimodal, corresponding to bursts (short ISIs) and isolated spikes (long ISIs).

#### Model optimization and parameter fitting

Parameter fitting of the computational models to experimental data was performed in Python using the Optuna optimization framework (Akiba et al., 2019). We used the sum of mean squared errors (MSE) between model and experimentally obtained features as the multi-objective loss function. Two distinct sets of features were considered separately: (i) action potential waveform features extracted from spike shape analyses and (ii) spike-train metrics including coefficient of variation, ISI mean and median, mean firing rate, burst fraction, and standard deviation of ISIs. Model parameters were initially constrained within ±20% deviation ranges from previously published parameter values (Destexhe et al., 1994; Li and Rinzel, 1994; Doiron et al., 2002; Toporikova and Chacron, 2009; Oprea et al., 2022). The optimization of these parameters was carried out using the non-dominated sorting genetic algorithm II (NSGA-II) which is a multi-objective optimizer (Deb et al., 2002) from Optuna, enabling efficient exploration of parameter space to minimize discrepancies between experimental and simulated neuronal behaviors.

#### Bifurcation and Sensitivity Analysis

We performed bifurcation analyses using AUTO (Doedel, 1981) to investigate dynamical transitions in the computational models. For the ELL pyramidal cell model, we conducted one-parameter bifurcation analysis with respect to the applied current (*I*_*app*_) and two-parameter bifurcation analyses with *I*_*app*_ paired with either the SK or NMDA maximal conductances (*g*_*SK*_, *g*_*NMDA*_, respectively) to explore the model behavior systematically.

For the modified HR model, we first executed bifurcation analysis over *I*_*app*_, followed by global sensitivity analysis using Sobol indices (Sobol, 2001; Saltelli et al., 2008) to quantify the influence of individual parameters on firing patterns. To do so, we classified simulations obtained during parameter optimization across all ELL pyramidal cells in our dataset (*n* = 20) as “successful” if the total loss (e.g. the error measure between model output and experimental data) is below a threshold of 10^−1^ (*L* < 0.1). Simulations exceeding this threshold (*L* ≥ 0.1) were excluded to ensure the analysis focused on parameter sets that yielded biologically plausible dynamics. For the retained successful simulations, we computed the total variance (*var*_*total*_) in the loss metric and calculated the Sobol indices (*S*_*i*_) using the equation:

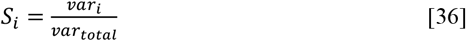

where *var*_*i*_ represents the variance attributable to parameter *i*. This approach allowed us to quantify the relative contribution of each parameter to the observed firing patterns.

#### Software and computational resources

All data analyses, simulations, feature extraction procedures, and parameter optimizations were performed using custom-written Python scripts executed on Compute Canada Alliance clusters. Bifurcation analyses were performed in AUTO-07 (Doedel et al., 2007), with a custom script for visualization of results. Model simulations were carried out using the Euler-Maruyama method (Kloeden et al., 1992) as the integration solver of the stochastic models. The excitatory and inhibitory pre-synaptic inputs in the biophysical ELL pyramidal cell model were generated using the algorithm described previously (Timmer and Koenig, 1995). For the biophysical model, all simulations were run using an integration timestep of 0.1 ms (100 kHz sampling rate).

The resulting membrane voltage traces were subsequently downsampled to 10 kHz to facilitate comparison with experimental recordings. This fine integration timestep was essential to ensure sufficient data points for accurate action potential shape analysis. For the modified HR model, simulations were performed at a 20 kHz sampling rate, after which the data were downsampled to 10 kHz to maintain consistency with experimental recordings.

## Authors Contribution

AA, MJC, and AK designed the research. MGM gathered the data. AA, MGM, analyzed the data. AA contributed to the computational modeling. AA, MJC, and AK wrote the initial draft, and all authors reviewed and revised the manuscript. MJC and AK supervised and acquired the financial support for this project.

## Acknowledgments

This research was funded by the Fonds de Recherche du Québec – Nature et Technologies (FRQNT) grant to MJC and AK, a Canadian Institutes of Health Research (CIHR) grant to MJC, and Natural Sciences and Engineering Research Council of Canada (NSERC) discovery grants to MJC and AK. AA was supported by the NSERC-CREATE in Complex Dynamics, Healthy Brains Healthy Lives (HBHL), Canada First Research Excellence Fund and the FRQNT fellowships.

## Declaration of Interests

The authors declare no competing interests.

## Notes

### Competing Interest Statement

The authors have declared no competing interest.

