## Supplementary Figures for "*In vivo* neural activity of electrosensory pyramidal cells: Biophysical characterization and phenomenological modeling"

### Supporting Information for
### *In vivo* neural activity of electrosensory pyramidal cells: Biophysical characterization and phenomenological modeling

Amin Akhshi[1], Michael G. Metzen[1], Maurice J. Chacron[1], Anmar Khadra[1*]

[1] Department of Physiology, McGill University, Montreal, QC, H3G 1Y6, Canada.

* Corresponding author: Anmar Khadra

#### Supplementary Figures

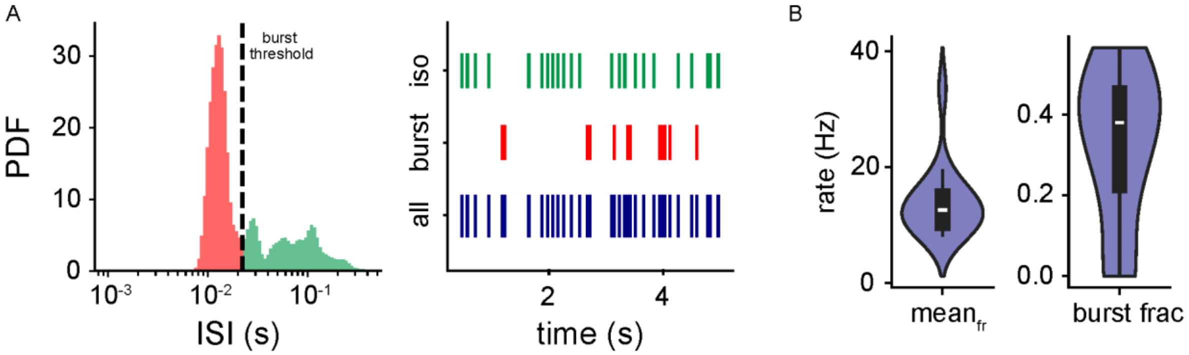


**Supplementary Figure 1:** Quantification of *in vivo* burst firing statistics in ELL pyramidal cells. **(A)** Illustration of burst definition and spike train separation for a representative neuron. Left: The burst threshold (dashed line) is set at the local minimum, separating short ISI mode (red) from longer ISIs associated with isolated spikes (green). Right: Spike raster plot of the same cell showing all recorded spikes (dark blue), and the resulting separation into burst spikes (red) and isolated spikes (green) based on the ISI threshold. **(B)** Distribution of mean firing rates (left) and burst fractions (right) all for recorded pyramidal cells ($N = 3$ fish, $n = 20$ cells).

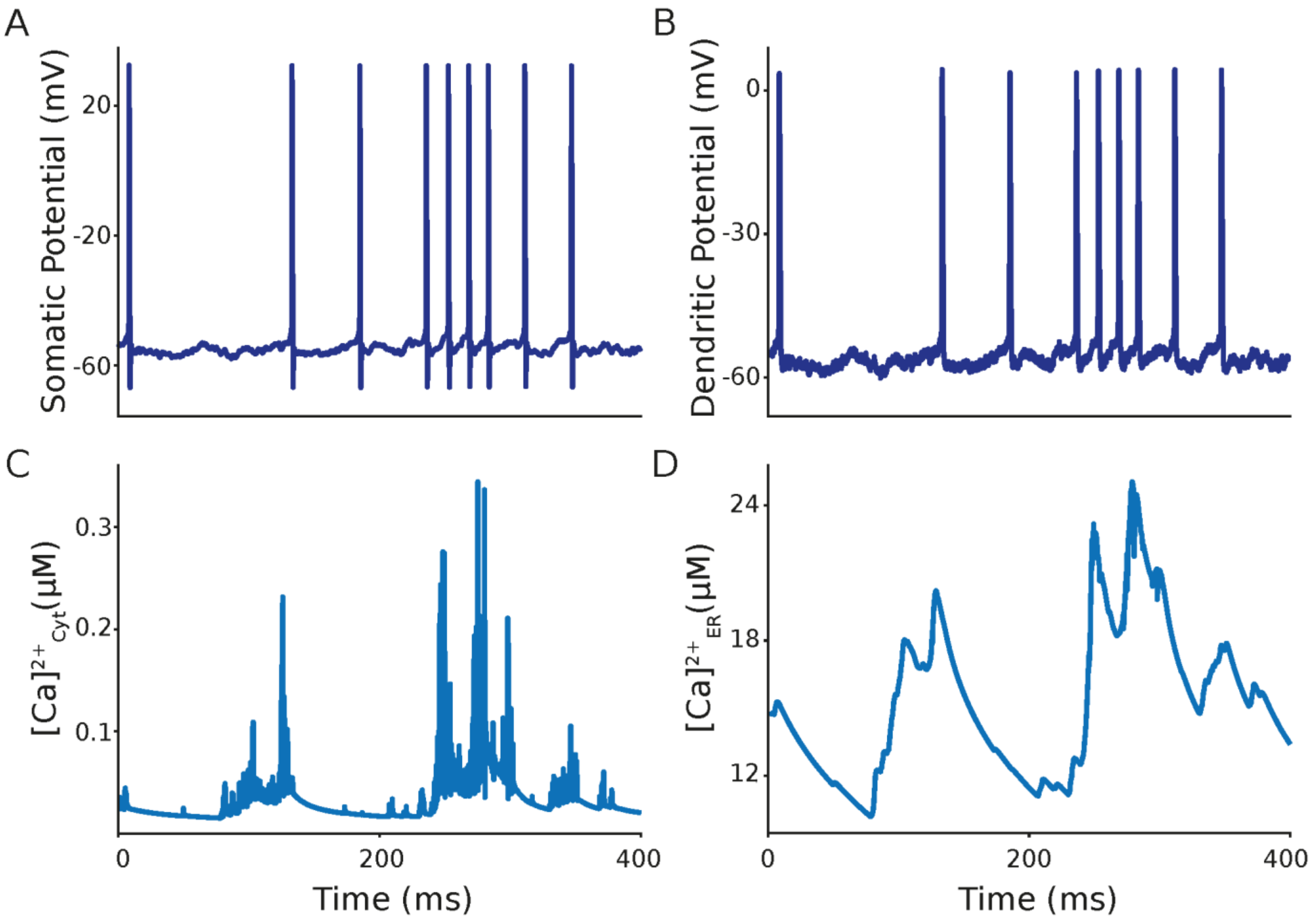


**Supplementary Figure 2:** ***Somatic and dendritic voltages and intracellular $Ca^{2+}$ dynamics.*** **(A, B)** Somatic (A) and dendritic (B) membrane voltages. The somatic action potential propagation to the dendritic compartment generates dendritic action potentials upon activation of $Na^{+}$ channel in the dendrite which in turn backpropagates to the soma through electrodiffusion coupling. Spike train traces show a gradual decrease in consecutive ISIs during the burst period but no evidence of the depolarizing afterpotential (DAP) growth or failure in the dendritic action potentials upon the termination of the burst similar to *in vivo* recordings. **(C, D)** Time series of the $Ca^{2+}$ concentration within the cytosol (C) and the ER (D) from dendritic compartment. $Ca^{2+}$ influx into the dendritic compartment through NMDA receptors or $Ca^{2+}$ release from the ER activates SK channel which in turn promotes afterhyperpolarization in the membrane potential. This effect is the main reason for DAP termination and elimination of dendritic failure, causing the difference in the burst mechanism between *in vitro* and *in vivo* recordings.

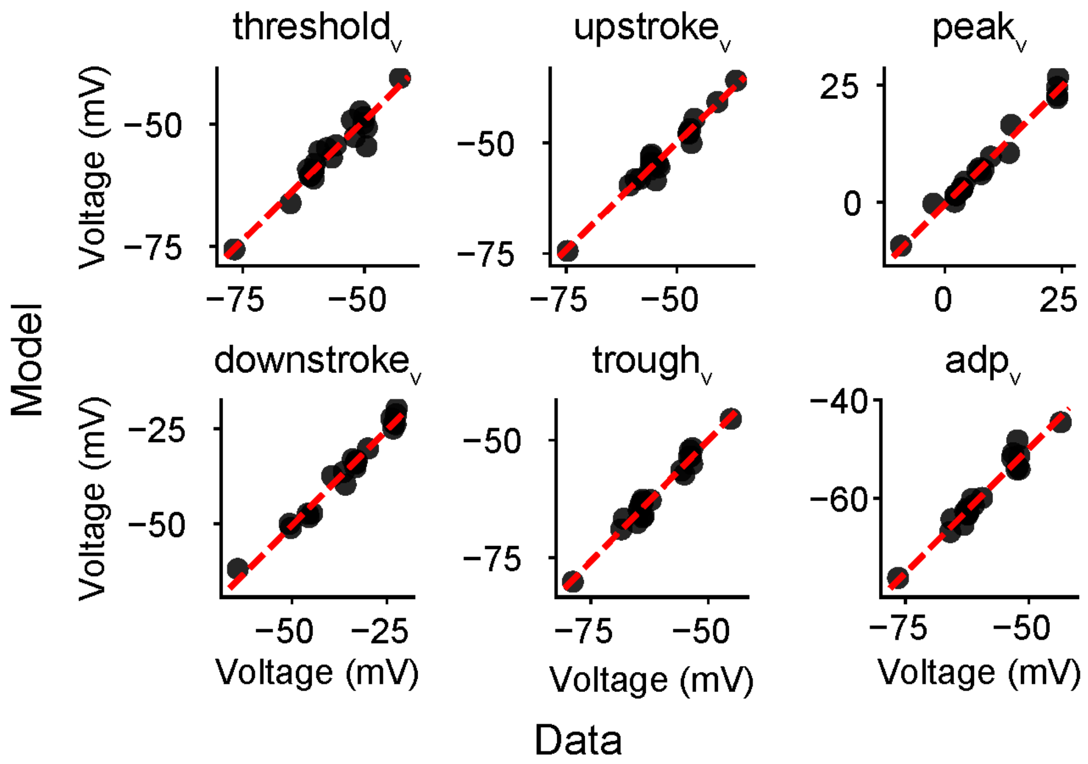


**Supplementary Figure 3:** ***Comparison of action potential features between experimental data and biophysical model simulations across the population***. Scatter plots comparing the mean value of each action potential feature between experimental data (data) and corresponding fitted model simulations (model) for all recorded ELL pyramidal cells (n=20). Features shown from left to right : spike threshold (threshold$_v$), midpoint of the upstroke phase (upstroke$_v$), peak amplitudes (peak$_v$), midpoint of the downstroke phase (downstroke$_v$), trough amplitudes (rough$_v$) and amplitude of afterdepolarization potential (adp$_v$). Points cluster closely around the identity line, indicating strong agreement. (linear fit: $r_{threshold}=0.96$, $p_{threshold}=4.8\times10{-}4$; $r_{upstroke}=0.98$, $p_{upstroke}=3.4\times10{-}5$; $r_{peak}=0.99$, $p_{peak}=5.2\times10{-}4$; $r_{downstroke}=0.99$, $p_{downstroke}=3.5\times10{-}4$; $r_{trough}=0.98$,$p_{trough}=8.1\times10{-}5$; $r_{adp}=0.98$, $p_{adp}=1.53\times10{-}8$).

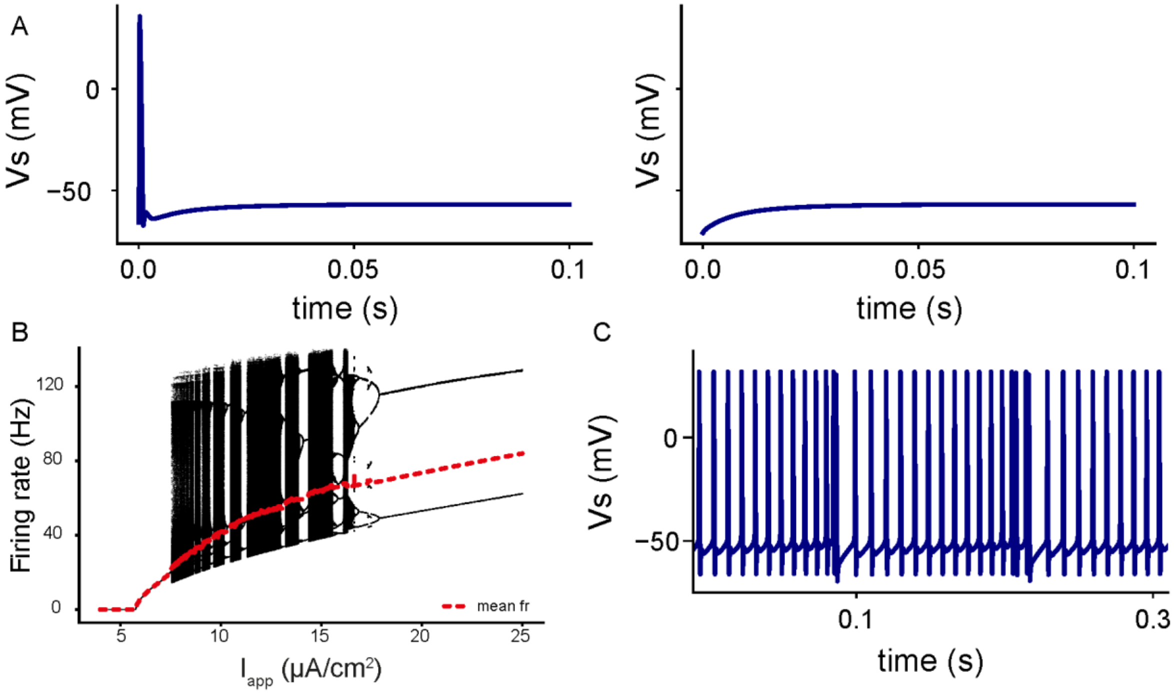


**Supplementary Figure 4: (A)** Membrane potential simulations of the biophysical model (0.1 s duration) with identical parameter values but different initial conditions, using a subthreshold value of the depolarizing current ($I_{app}$) below the SN1 bifurcation point (see Fig. 4A). Depending on the initial condition of the system, the model either returns directly to the resting state (right panel) or generates a single transient spike before returning to rest (left panel). **(B)** The firing rate of the biophysical two compartment model detected within a time range of 20 seconds with respect to $I_{app}$, showcasing the chaotic dynamics exhibited by this model. Red curve represents the average firing rate. (C) Simulation of the biophysical model after blocking SK and NMDA currents ($g_{SK} = 0$, $g_{NMDA} = 0$). In the absence of these currents, the model exhibits prolonged bursts with prominent depolarizing afterpotentials (DAPs), reverting to dynamics characteristic of the original *in vitro* ghostbursting model.

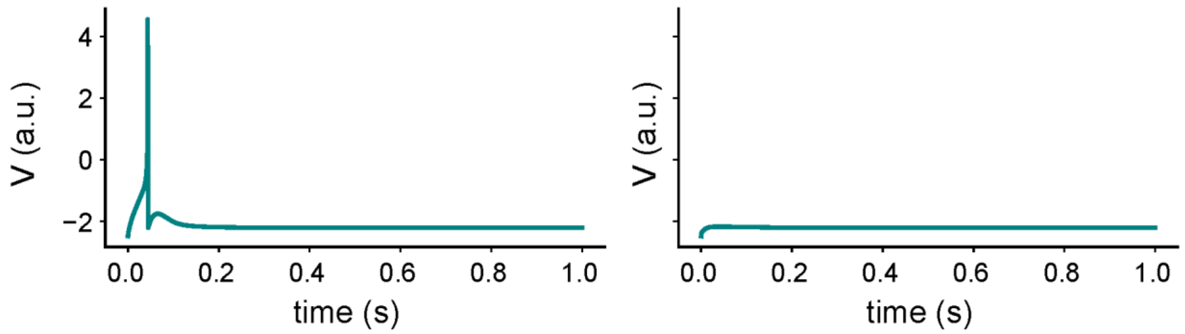


**Supplementary Figure 5:** ***Initial condition dependence of subthreshold responses in the modified HR model***. Membrane potential simulations (1s duration) for the same parameter values using two different initial conditions with the applied current $I_{app}$ below the spiking threshold. Depending on the specific initial condition, the model either generates a single transient spike before returning to rest (left) or returns directly to the resting state without spiking (right).

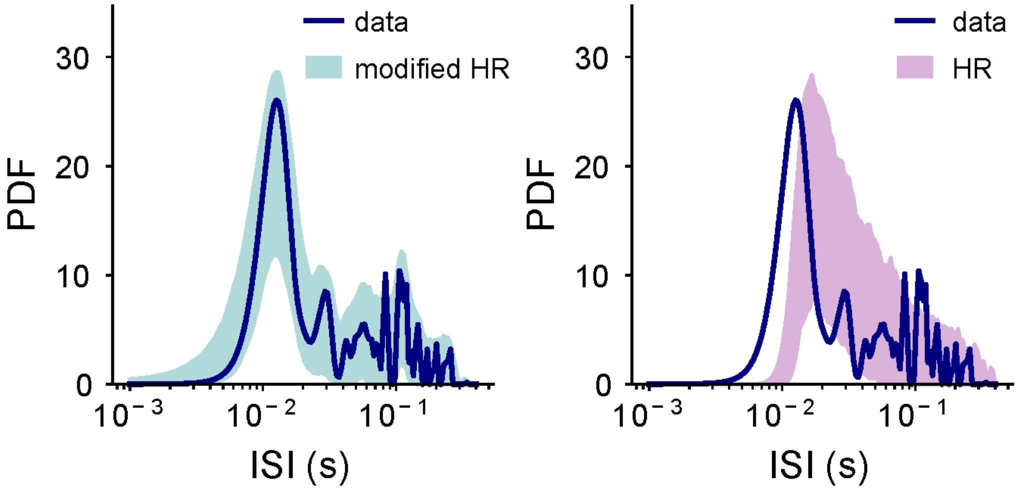


**Supplementary Figure 6:** Comparison of the interspike interval (ISI) distributions between an example recorded ELL pyramidal cell (blue) and ISIs from model simulations of modified HR model (teal, left) and classic HR model (purple, right) both having same parameter values and stochastic synaptic input (Kolmogorov-Smirnov test vs data: $D_{\text{modified HR}}= 0.146$, $p_{\text{modified HR}}=0.778$; $D_{HR}=0.383$, $p_{HR}= 0.0065$).
